# One Sea, Different Whales: Genomics Sheds Light on a Small Population of Fin Whales

**DOI:** 10.1101/2025.06.16.659691

**Authors:** Roberto Biello, Alessio Iannucci, Silvia Fuselli, Elisa Desiato, Jorge Urban R., Maria Cristina Fossi, Cristina Panti, Annalaura Mancia

## Abstract

Whales play a crucial role in marine ecosystems, yet many populations face growing environmental pressures. Among them, the fin whale, *Balaenoptera physalus*, from the Mediterranean Sea remains poorly characterized at the genomic level despite its classification as Endangered and decreasing in population size due to habitat degradation, climate change, and human-induced disturbances. Previous studies based on bioacoustics and telemetry data suggest the presence of resident and migratory subgroups, but the genetic isolation of this population remains uncertain. Here, we sequenced and analysed for the first time whole genomes of the Mediterranean fin whale to provide a comprehensive genomic characterization of this population, assessing its genomic variability, genetic load, population structure and potential for adaptation to environmental stressors. By comparing genomic data from Mediterranean fin whales with populations from the North Atlantic, North Pacific, and the Sea of Cortez (also sequenced in the present study), we aim to determine the degree of genetic isolation and contextualize the Mediterranean population within a broader evolutionary and conservation framework. Our results showed that while Mediterranean fin whales form a distinct genetic cluster, they are not entirely isolated from North Atlantic populations, with gene flow persisting at low levels. Furthermore, we identified a complex substructure within the Mediterranean, supporting the existence of a resident subpopulation. Although this population preserves moderate levels of genomic diversity and adaptive potential, it remains vulnerable to genomic erosion due to continued demographic decline, limited connectivity, and increasing environmental stressors. These findings underscore the urgency of targeted conservation actions and long-term genetic monitoring, especially as climate change accelerates and introduces increasingly unpredictable selective pressures that may exacerbate the risk of genomic erosion and threaten population viability.

## 1 INTRODUCTION

Whales have long captured human fascination, not only due to their massive size but also because of their complex behaviours, migratory patterns, and ecological importance. Among them, baleen whales play a crucial role in maintaining ocean ecosystems by facilitating nutrient cycling and influencing marine food webs (Ganley et al., 2022; Durfort et al., 2022). However, many species face increasing environmental pressures, from climate change to human-induced disturbances, making their study more critical than ever (Peters et al., 2022; Fossi et al., 2020; Panti et al., 2019; Tulloch et al., 2019).

As the climate crisis accelerates, genomic studies have become powerful tools for investigating population structure, evolutionary history, and patterns of isolation. By analysing genetic differentiation, we can determine whether a population is truly isolated or maintains some connectivity with neighbouring groups. This is essential for conservation efforts, as isolated populations with low genetic diversity may be more vulnerable to environmental changes, disease, pollution, and habitat degradation (Cabrera et al., 2022; Simmonds & Eliott, 2009). Furthermore, understanding the degree of isolation can clarify the evolutionary processes shaping populations, providing insights into how adaptation occurs across different ecological contexts (Pallin et al., 2023; Clapham et al., 1999).

The fin whale (*Balaenoptera physalus*) is the second-largest whale species and mainly inhabits temperate and polar latitudes, with an apparent equatorial hiatus separating Northern and Southern Hemisphere populations (Edwards et al., 2015; Aguilar & García-Vernet, 2018). Like most mysticetes, fin whales typically undertake long-range annual migrations with a seasonal cycle between high-latitude feeding grounds and low-latitude breeding and calving grounds in winter, where feeding is absent (Kellogg, 1929; Katona & Whitehead, 1981). As an exception to these migration patterns, resident populations of fin whales such as those in the Sea of Cortez and the Mediterranean Sea have also been documented (Tershy et al., 1993; Bérubé et al., 1998; Forney & Barlow, 1998; Bérubé et al., 2002; Palsbøll et al., 2004; Urban et al. 2005; Stafford et al., 2007; Moore et al. 2006; Mizroch et al. 2009). The Mediterranean fin whale population has been identified as distinct from the North Atlantic populations through bioacoustics and telemetry studies (Bentaleb et al., 2011; Cotté et al., 2011; Castellote et al., 2012; Panigada et al., 2017). Moreover, previous studies suggested the presence of both resident and migratory subgroups within the Mediterranean Sea, adding complexity to its structure (Castellote et al., 2012). To date, the genetic characterization of this fin whale population remains poor, with only a few studies based on traditional molecular markers conducted to determine whether this population is genetically isolated (Bérubé et al., 1998; Palsbøll et al., 2004).

Mediterranean fin whales aggregate during the summer months on the feeding grounds of the Pelagos Sanctuary for Marine Mammals, located in the northwestern Mediterranean Sea (Notarbartolo di Sciara et al., 2003) and presumably migrate to the southern Mediterranean Sea during winter (Panigada et al., 2011). This area is characterized by high offshore primary productivity, which attracts a variety of predators, including eight cetacean species and many large marine vertebrates (Notarbartolo di Sciara et al., 1993, Coll et al., 2012). This remarkable biodiversity coexists with extremely high human pressure and threats, including habitat degradation, climate change, heavy maritime traffic, anthropogenic noise and chemical pollution (Espada et al., 2024; Fossi et al., 2016) along with large amounts of plastic debris and microplastics (Collignon et al., 2012; Fossi et al., 2013; Cózar et al., 2015). These threats, combined with their restricted distribution within the semi-enclosed and heavily impacted Mediterranean basin and an inferred continuing decline in the number of mature individuals, recently estimated at around 2,500, have led to their classification as Endangered on the IUCN Red List of Threatened Species (Panigada et al., 2021).

Given the limited genomic knowledge and threats faced by fin whales, this study provides the first genomic characterization of the Mediterranean fin whale population, aiming to more accurately reconstruct its population structure, as well as its demographic and evolutionary history. In particular, we infer the potential for adaptation to increasing environmental disturbances by using the estimated level of genomic variability within the Mediterranean population and genetic load as proxies. Furthermore, comparison with data available in the literature, as well as new data generated in this study from populations of the same species in different geographical areas, allowed us to: (i) assess the degree of isolation of the Mediterranean population from North Atlantic populations, with which some level of gene flow is hypothesized, and (ii) contextualize Mediterranean fin whale genomes within a broad geographical perspective, including North Pacific populations known for their relatively large population sizes. Additionally, the Sea of Cortez population was included, as it provides an important point of comparison with a nearly completely isolated and small-sized population.

## 2 MATERIAL AND METHODS

### 2.1 Genome scaffolding and annotation

We scaffolded the *B. physalus* genome (Wolf et al., 2022) using sequence data from DNA ZOO (https://www.dnazoo.org/assemblies/balaenoptera_physalus). These data include in vivo Hi-C data (SRR16970344). Chromap v0.2.4 (Zhang et al., 2021) was used to align the HiC reads to the genome using parameters for HiC data (--preset hic) and remove pcr duplicates (--remove-pcr-duplicates). Output .sam file was converted to a sorted .*bam* file with samtools v1.11 (Li et al., 2009). The latter along with the reference assembly served as the input files for scaffolding with YaHS v1.2 (Zhou et al., 2023). This was implemented using standard option, and YaHS outputs were converted using the “juicer_pre” function of YaHS to Juicebox Assembly Tools (jbat) (Dudchenko et al., 2018) compatible files for manual curation visually within Juicebox. Following manual curation, edits were applied to the scaffold assembly using “juicer_post”. We assessed the quality of the genome assembly by searching for conserved, single copy, Mammalia genes (n = 9,226) with Benchmarking Universal Single-Copy Orthologs (BUSCO) v5.3.2 (Simão et al., 2015) and by analysis of k-mer spectra with MERQURY (Rhie et al., 2020) to compare k-mer content of the raw sequencing reads to the k-mer content of the assembly.

Using the homology-based analysis, we identified the known transposable elements (TE) within the *B. physalus* genome using Repeatmasker v4.0.7 (Smit et al., 2013) with the combined database of RepBase (Jurka et al., 2005) and Dfam Consensus (Wheeler et al., 2013). Repeatmasker was launched with options “-e ncbi-species mammalia-xsmall-gff” (Jurka et al., 2005).

For gene prediction, we first downloaded RNA-seq reads available on NCBI from various tissues of closely related species (**Table S1**). Quality control and trimming for adapters and low-quality bases (quality score <20) of the raw reads were performed using fastqc v0.11.8 (Andrews, 2010) and TrimGalore v0.5.0 (https://github.com/FelixKrueger/TrimGalore), respectively. High-quality reads were then mapped to the soft-masked assembly with HISAT2 v2.2.1 (Kim et al. 2015) and sorted with samtools v1.11 (Li et al., 2009). All the BAM files were filtered to remove invalid splice junctions with Portcullis v1.2.4 (Mapleson et al. 2018). Filtered RNA-seq alignments were passed to Braker v3 (Hoff et al., 2016, 2019; Brůna et al., 2021; Gabriel et al., 2023), together with protein sequences of nine closely related species from the order Artiodactyla including two *Balaenoptera* species (**Table S2**). The Braker gene prediction pipeline was run with the options “--softmasking”. This pipeline uses StringTie (Pertea et al., 2015) to assemble the RNA-seq reads followed by rounds of GeneMark and AUGUSTUS training and gene prediction (Stanke and Waack 2003). Gene sets were combined with TSEBRA (Gabriel et al., 2021). The completeness of the final gene set was checked with BUSCO v5.3.2 (Simão et al., 2015) using the longest transcript of each gene as the representative transcript.

Sequences were searched against the nonredundant NCBI protein database using DIAMOND v0.9.10 (Buchfink et al. 2015) with an E-value cut-off of ≤ 1×10^-5^. BLAST2GO v5.0 (Conesa et al. 2005) and INTERPROSCAN v2.5.0 (Quevillon et al. 2005) were used to assign Gene Ontology (GO) terms. Protein domains were annotated by searching against the InterPro v32.0 (Hunter et al. 2012) and Pfam v27.0 (Punta et al. 2012) databases, using INTERPROSCAN v5.52 (Quevillon et al. 2005) and HMMER v3.3 (Finn et al. 2011), respectively.

### 2.2 Phylogeny

Orthologous groups in Cetacea genomes were identified from the predicted protein sequences of *B. physalus* and 11 other Cetacea genomes already published (see **Table S3**). As outgroups, we included the genomes of two Artyodactyla: *Camelus dromedarius* (GCF_036321535.1) and *Hippopotamus amphibius kiboko* (GCF_030028045.1) (**Table S3**). We used the longest transcript to represent the gene model when several transcripts of a gene were annotated. Orthofinder v2.5.4 (Emms & Kelly, 2019) with diamond v0.9.14 (Buchfink et al. 2015), Multiple Alignment using Fast Fourier Transform (MAFFT) v7.305 (Katoh & Standley, 2013) and with RAxML v8.2.12 (Stamatakis 2014) were used to cluster proteins into orthogroups, reconstruct gene trees and estimate the species tree.

### 2.3 Sampling, DNA isolation and sequencing

A total of 13 tissue samples were collected as skin biopsies from individual free-ranging fin whales in the Mediterranean Sea (MED) in 2018-2019 (**Table S4**). Samples were collected by remote dart-sampling in accordance with national and international regulations and under sampling permits n. 0018799/PNM released to the University of Siena from the Italian Ministry of Environment and Energy Safety and the Italian National Institute for Environmental Protection and Research (ISPRA). The research also permits included the necessary ethical approval for sample collection, analysis and use. Additional individuals sampled in the Sea of Cortez were sequenced in this study (SOC, N = 7; **Table S4**). Skin tissues obtained from darts were snap-frozen in liquid nitrogen and then stored at −80°C until processing. DNA was extracted from 20 mg of skin tissue using the Wizard® Genomic DNA Purification Kit (Promega) following the manufacturer’s instructions. DNA integrity was assessed by electrophoresis on a 1% agarose gel and DNA concentration was measured by fluorometric analysis using the Qubit™ 4 Fluorometer (Invitrogen, Waltham, USA). Short-read genomic libraries were constructed using a Illumina DNA PCR-Free Prep Kit (Illumina) according to the manufacturer’s protocol. For sequencing, libraries were run paired-end 2×150 bp using an Illumina NovaSeq 6000 System (Illumina, Inc. San Diego, CA, USA) on a S1 flow cell with 500 Gb as final output.

### 2.4 Mapping

Demultiplexing and conversion of sequencing data from bcl to fastq formats were performed using bcl2fastq v2.20 (Illumina). Quality control of the reads was assessed with FastQC v0.11.8 (Andrews, 2010). Reads were then processed with AdapterRemoval v2 (Schubert, Lindgreen, & Orlando, 2016) to remove residual Illumina adapters. Read tails with a mean Phred-quality score <10 over a 4 bp sliding window were trimmed and subsequently aligned to the *B. physalus* reference genome produced in this study using the mem algorithm implemented in the bwa v0.7.15 aligner (Li & Durbin, 2009). Alignments in sam format were sorted, indexed and compressed in bam format using samtools v1.9 (Li & Durbin, 2009). PCR duplicates, produced during library preparation, and optical duplicates were removed using the MarkDuplicates tool in the Picard Toolkit v2.18.20 (http://broadinstitute.github.io/picard/). Regions close to indels showing putative alignment errors were identified and realigned using the RealignerTargetCreator and the IndelRealignment tools in GATK v3.5 (McKenna et al., 2010). Alignment statistics were calculated using the CollectAlignmentSummaryMetrics tool, and bam files were validated with the ValidateSamFile tool of the Picard Toolkit v2.18.20. Observed coverage was computed using the depth command of samtools v1.9 with the “-aa” flag activated.

We also downloaded paired-end reads of 27 additional samples (see **Table S5**) of *B. physalus* sampled from Iceland (ICE, N = 11), Svalbard archipelago (SVA, N = 7), North Pacific (NPA, N = 9) and 5 additional SOC individuals for which we applied the same informatics pipeline as described above.

### 2.5 Mitochondrial DNA

Raw reads of the 20 individuals sequenced in this work were aligned to the fin whale mitogenome reference sequence (Árnason et al. (1991), NCBI access no. NC001321) as described in paragraph 2.3. Consensus sequences were created with angsd v0.932 using options -minQ 20 -minMapQ 30 - setMinDepth 3. Additional complete mitogenome sequences were retrieved from the literature (**Table S6**) for a total of 265 sequences. Sequences were aligned using Geneious Prime 2020.1.1 (Kearse et al., 2012), visually checked for accuracy and trimmed to the shortest available sequence (16,423 bp).

All 265 sequences were used to build an mtDNA phylogenetic tree using RAxML v8.2.7 implemented in Geneious Prime 2020.1.1 by applying the GTR GAMMA nucleotide model with rapid bootstrapping and search for best-scoring maximum-likelihood trees across 100 bootstrap replicates (Stamatakis, 2014). We used the complete mtDNA sequence of the *Balaenoptera musculus* outgroup (NCBI accession no.: NC001601). Figtree v1.4.4 was used to visualise and edit the phylogenetic tree (http://tree.bio.ed.ac.uk/software/figtree).

### 2.6 Relatedness

To detect the presence of close relatives we estimated coefficients of relatedness with ngsRelate (Hanghøj et al., 2019), which requires the use of genotype likelihoods and population allele frequencies. The coefficient of relatedness estimator used in ngsRelate is given by considering all possible patterns of identity-by-descent sharing between two individuals to account for the possibility of the individuals being inbred (Hedrick & Lacy, 2015). Inferred kinship coefficients suggested that couples MED01-MED02, MED10-MED12 (MED) and BP1402-BP1602 (SVA) were siblings or parent–offspring pairs (**Table S7**). Thus, among these pairs we retained the sample with higher coverage (MED02, MED10 and BP1402) for the following analyses.

### 2.7 Genotype likelihood and SNP calling

Genotype likelihoods were calculated on the 49 individuals using ANGSD (v0.933) (Korneliussen et al., 2014) with the GATK model (“-GL 2”) and the following parameters: “-doMajorMinor 1 - minMapQ 20 -minQ 20 -doMaf 1 -SNP_pval 1e-3 -minMaf 0.05 -doGlf 1 -minInd 30”. We masked genomic regions from the reference containing repetitive elements (as described in Genome scaffolding and annotation) and having a low mappability score (p < 1) computed using gem (Derrien et al., 2012) by setting a maximum mismatch of 4% in a 150bp read. The same filters were used in subsequent analyses conducted with ANGSD, unless specified otherwise.

For analyses where a higher coverage is recommended, we used BCFtools v1.11 (Danecek et al. 2021) mpileup and call function identify SNPs in each sample. VCFtools v.0.1.16 (Danecek et al. 2011) was used to filter the dataset to only retain biallelic SNPs with SNP quality scores (QUAL) ≥30 with the option “--remove-indels --max-alleles 2 --max-missing 0.9 --minQ 30 --min-meanDP 5 --max-meanDP 35 --minDP 5 --maxDP 35”. Finally, all sites included in repeated genomic regions or in regions with a low mappability score, calculated using the GEM algorithm (Derrien et al., 2012) and setting a maximum mismatch of 0.04 in a 150 bp read, were removed.

### 2.8 Population structure

We used the genotype likelihoods of autosomal chromosomes to perform a number of structure analyses. We performed a PCA using PCAngsd v0.96 (Meisner & Albrechtsen, 2018) both on the complete dataset and excluding individuals from the Sea of Cortez and North Pacific.

In order to infer admixture proportions and model-based individual clustering from genotype likelihoods, we ran NGSadmix (Skotte et al., 2013) on the complete dataset. The analysis was performed assuming from one to ten ancestral populations (K) and doing 100 independent runs in each case. In the cases where the admixture analysis converged the maximum-likelihood result was selected. Furthermore, we evaluated the admixture model fit at every converged K with evalAdmix (Garcia-Erill & Albrechtsen, 2020).

We evaluated genetic clustering based on genetic distances by building a phylogenetic tree based on an identity-by-state (IBS) matrix. We used the -doIBS 1 option in ANGSD (Korneliussen et al., 2014) to generate the IBS matrix of pairwise genetic distances, applying the same settings as listed in paragraph 2.3. The distance matrix was then converted to the nexus format using the phangorn R package (Schliep, 2011), and SPLITSTREE v4.14.6 (Huson & Bryant, 2006) was used to obtain a phylogenetic network according to the Neighbor-net algorithm (Bryant & Moulton, 2004).

We estimated F_ST_ indices between each population pair, extracting the F_ST_ values from the corresponding unfolded 2DSFS using Hudson’s estimator, which is less sensitive to differences in sample size between populations (Bhatia et al., 2013). Both mitochondrial and nuclear data assigned the individual SOC02 sampled in the Sea of Cortez, to the North Pacific cluster. Based on these results, this individual was removed from the F_ST_ analysis.

### 2.9 Genomic diversity

Genomic diversity of individuals and populations was evaluated using the Watterson’s Theta estimator (*θ_w_* Watterson, 1975). The number of segregating sites was counted for each callable region, defined by genomic intervals showing good mappability, low repetitiveness and appropriate coverage levels (see paragraph 2.3). The total number of segregating sites was first divided by the (n-1) harmonic number, where n is the number of haploid chromosome copies, and then by the total size of the callable regions to obtain the per-base estimator *θ*_W_. The same approach was used to obtain a *θ*_W_ estimate over neutral regions and exons. Neutral regions were defined by callable regions located in intergenic regions that were at least 25 Kb from the closest gene. Exon regions were extracted directly from the reference genome annotation by merging overlapping elements in different strands and discarding portions that were not in callable regions. Among these exons, exon 2 (270 bp) of one of the most polymorphic loci in vertebrate genomes, the *DQB-1* locus in the major histocompatibility complex (MHC), was chosen to represent a genomic region where high genetic variability is expected, as an indication of good adaptive potential against pathogens. MHC *DQB-1* exon 2 was analysed for 7 MED and 2 SOC individuals from this study, as well as for 9 NPA, 6 SVA, 11 ICE, and 5 SOC individuals from previous studies (see paragraph 2.3). Additionally, sequences from two closely related species, *Balaenoptera musculus* (MUS, N=5) and *Megaptera novaeangliae* (MEG, N=5), were obtained from public repositories (**Table S8**). Genetic diversity indices were calculated to assess variability as previously described.

### 2.10 Inbreeding

Runs of homozygosity (ROH) were first identified by estimating the heterozygosity levels in 100 Kb non-overlapping windows using Rohan’s probabilistic method (Renaud, Hanghøj, Korneliussen, Willerslev, & Orlando, 2019). Each genomic segment was then defined to be in ROH based on a Hidden Markov Models (HMM) classifier. The analysis was performed on bam alignments considering base and mapping errors. We used a transition/transversion rate of 2.019, estimated by vcftools across the entire dataset, and an expected *θ*_W_ in ROH regions (rohmu flag) of 5×10^-4^. The parameter *θ*_W_ was estimated by either including or excluding ROH regions. Despite the method being developed to provide reliable ROH estimates for different coverages (>5×) (Renaud et al., 2019), individuals sequenced at higher coverage were downsampled to 10× in order to facilitate comparison with the other samples. ROH regions ≥ 100Kb were used to estimate the fraction of the whole genome that was in ROH state (FROH). We then calculated the fraction of the genome in ROH obtained from the output file HMM posterior decoding using .mid estimates of heterozygosity, and we binned them by ROH size (ROH > 1 Mb).

### 2.11 Demographic history

Trajectories of effective population size (Ne) through time were inferred using a Multiple Sequentially Markovian Coalescent - MSMC (Schiffels & Wang, 2020) on one representative high-coverage sample for SOC, NPA, ICE and SVA populations and all samples with a coverage above 13x in MED population (**Table S4-S5**). Input data was generated using the “generate_multihetsep.py” script selecting all callable segments from 21 autosomal scaffolds. Effective population size and times were scaled using a mutation rate of 1.39 × 10−8 substitutions/site/generation and a generation time of 25.9 years (Árnason et al. 2018).

### 2.12 Genetic load

We first polarised each SNP as ancestral or derived using two outgroups: *Balaenoptera musculus* (5 individuals) and *Megaptera novaeangliae* (5 individuals) (**Table S8**). We defined the ancestral allele as the allele present in the two outgroups using a custom python script. All sites where at least one of these outgroups was heterozygous were discarded to maximise the confidence in the ancestral allele definition. We assessed the mutational load in through two distinct methods. We restricted these analyses to the protein coding sequences (CDS), extracting the CDS of the longest transcript of each gene of the annotation and merging the overlapping windows with bedtools v2.30 (Quinlan et al. 2010). First, we run SnpEff v5.0 (Cingolani et al. 2012) on the polarised SNPs present in CDS to classify each mutation as synonymous, missense (i.e., non-synonymous) or nonsense (including stop-gained, start-gained, start lost, stop codons, splice donor variant and splice acceptors). For each class of putatively deleterious mutation (missense and nonsense mutations), the genetic load was separated into two components: i) the masked load estimated as the individual number of heterozygous sites, and ii) the realized load estimated as the individual number of homozygous derived sites (Bertorelle et al. 2022). Secondly, we measured the relative mutational load in each individual as the number of derived alleles at sites that are under strict evolutionary constraints (i.e., highly conserved) and thus likely to be deleterious using Genomic Evolutionary Rate Profiling scores (GERP). GERP scores for the *B. musculus* in bigwig format was obtained from a multiple alignment with 91 mammal species downloaded from ENSEMBL (https://ftp.ensembl.org/pub/current_compara/conservation_scores/91_mammals.gerp_conservation_score/). We firstly aligned the *B. physalus* assembly to the *B. musculus* assembly using minimap2 with parameter -cx asm20. The GERP scores were subsequently transferred to *B. physalus* reference coordinates using Transanno between *B. musculus* and *B. physalus* genome. Our analysis considered both heterozygous positions (counted as one allele) and homozygous positions (counted as two alleles), recognizing that the mutational impact of heterozygous positions involves additional assumptions regarding the dominance coefficient. GERP identifies constrained elements within multiple alignments by quantifying substitution deficits, reflecting substitutions that would have occurred if the element was neutral DNA but did not due to functional constraints, accounting for phylogenetic divergence. The individual relative mutational load was calculated by summing the number of derived alleles above a GERP score of four (highly deleterious).

## 3 RESULTS

### 3.1 The fin whale reference genome

The previous versions of the fin whale (*Balaenoptera physalus*) genome assemblies (Yim et al., 2014; Wolf et al., 2022; DNA Zoo) had a total length between 2.4 and 2.7 Gb (**Table 1**). Using the HiC reads (DNA Zoo; SRR16970344), we scaffolded the fin whale genome (Wolf et al., 2022) which resulted in a final assembly of 12,892 scaffolds, and an N50 and L50 of 109.1 Mb and 9 scaffolds, respectively (**Table 1**). After manual curation, the final assembly comprised 2.410 Gb, with 96.9% of the assembly anchored into 22 chromosomes (21 autosomes + X chromosome) (**Figure 1a**, **Figure S1**, **Table 1**), consistent with the fin whale 2n karyotype being 42 chromosomes + XY chromosomes (Arnason 1969). Based on coverage analyses, we discovered the sex chromosome in the genome assembly (**Figure S2**), showing 50% lower coverage than the autosomes. The lengths of the 22 chromosomes ranged from 182 to 35 Mb. The final assembly has a BUSCO completeness score of 93.5% using the Mammalia gene set, a per-base quality (QV) of 29.05 and a k-mer completeness of 91.62. Thus, the new assembly (Bphy_ph2.v2) represents a near complete and highly contiguous assembly and a significant improvement on the existing assemblies of this species (**Table 1**). The identification of repetitive elements resulted in a 38.2% repeat content, falling within the range of repeat contents for other Cetacean species (Tollis et al., 2019; Fan et al., 2019). In total, 25,321 protein-coding genes were predicted. The BUSCO completeness of the gene annotation using the same Mammalia gene set was 89.8%. To place the genome assembly in a phylogenetic context, we compared the proteome of *B. physalus* (which includes the complete set of annotated protein-coding genes) to those of 12 other cetacean species with fully sequenced genomes (see **Table S3**). Using a maximum likelihood approach, we conducted a phylogenetic analysis based on a concatenated alignment of 5,464 conserved single-copy genes. The resulting species tree was fully resolved, with 100% support at all nodes (**Figure 1b**). The tree confirmed the closer phylogenetic relationship of *B. physalus* with the humpback whale (*Megaptera novaeangliae*) compared to other *Balaenoptera* species.

**Figure 1.**
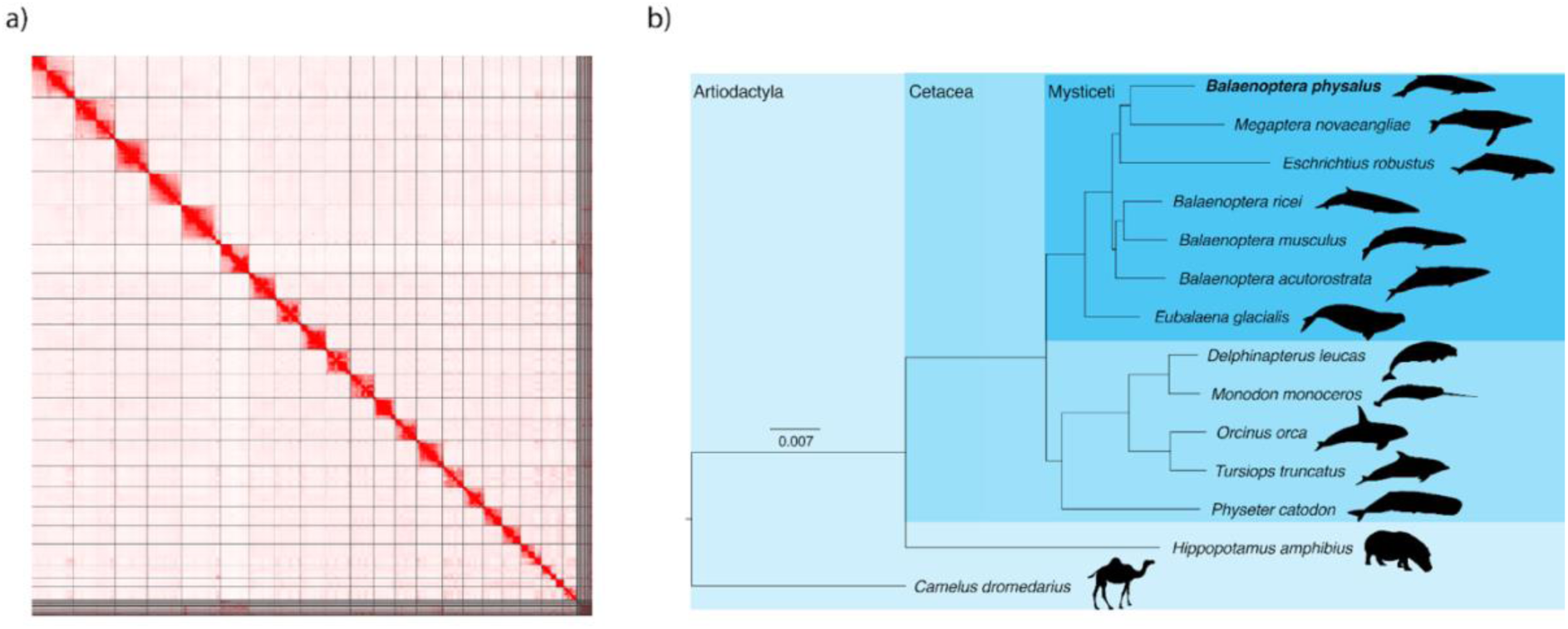
The fin whale reference genome. a) HiC heat-map of genomic interactions. Interactions between two locations are depicted by a red pixel. Black lines depict scaffold boundaries for the 22 chromosome-length scaffolds. b) Maximum likelihood phylogeny of the fin whale and 11 other Cetacea species based on a concatenated alignment of 5,464 conserved one-to-one orthologues. The tree is rooted with *Hippopotamus amphibius* and *Camelus dromedarius*. Branch lengths are in amino acid substitutions per site.

**Table 1.** Metrics of the fin whale genome assembly and comparison with publicly available fin whale genome assemblies.

| Genome Assembly | *Baphy* | *SBiKF_Bphy_ph2* | *Balaenoptera_physalus_HiC* | *Bphy_ph2.v2* |
| --- | --- | --- | --- | --- |
| Reference | Yim et al., 2014 | Wolf et al., 2022 | DNA Zoo | This study |
| GenBank/ENA accession # | GCA_008795845.1 | GCA_023338255.1 | / | / |
| Base pairs (Gb) | 2.462 | 2.41 | 2.754 | 2.41 |
| Total scaffolds | 62,302 | 13,140 | 1,361,899 | 12,892 |
| Scaffold N50 (Mb) | 0.9 | 24.9 | 77.5 | 109.1 |
| Scaffold L50 | 803 | 27 | 13 | 9 |
| Longest scaffold (Mb) | 6.8 | 91.4 | 156.9 | 182.7 |
| Assembly in chromosomes | 0.00% | 0.00% | 72.70% | 96.90% |

### 3.2 Genotype calling

For each sample, the number of reads, percentage of aligned reads, and final mean coverage are provided in **Table S4-S5**. After filtering for mapping quality, scaffold size, sex-linked scaffolds, repetitive regions, and mappability, a total of 7,217,348 genomic sites were retained for downstream analyses, including those with low-depth data. For analyses requiring high-depth data, a subset of samples with at least 10× coverage was used, resulting in 5,850,219 sites.

### 3.3 Population structure

The phylogenetic structure estimated from the complete mitogenomic haplotypes revealed three main clades that are strongly supported by bootstrap values and associated with distinct ocean basins (**Figure S3**). Cluster 1 primarily included haplotypes from the North Pacific (NPA) and the Sea of Cortez (SOC). Cluster 2 consisted of haplotypes from the Southern Hemisphere, along with two smaller clusters from the North Atlantic and NPA. Cluster 3 mainly included samples from the North Atlantic and the Mediterranean Sea (**Figure S3**). These results are consistent with those obtained in previous studies (Archer et al., 2013; Cabrera et al., 2019; Buss et al., 2023).

The analysis of nuclear DNA using Principal Component Analysis (PCA) highlighted distinct patterns of genetic differentiation across populations. Along the first principal component (PC1), there was a pronounced separation between Pacific and Atlantic populations, underscoring significant divergence between these two oceanic regions. Further examination within the Pacific populations revealed additional structure along PC2, where the isolated population from the SOC was clearly distinct from the NPA samples (Figure 2a). Focusing on the Atlantic and MED populations, PCA showed a slighter degree of differentiation. Among the Iceland (ICE), Svalbard (SVA), and Mediterranean (MED) groups, genetic divergence was indicative of an underlying population structure. Within this grouping, the MED samples stand out by exhibiting the highest level of internal genetic variation, which could reflect historical admixture with the close population from the Atlantic Ocean. The admixture analysis further complements these observations, revealing a similar genetic structure. At K=2, the most strongly supported number of genetic clusters (**Figure S4**), there was a clear distinction between Atlantic and Pacific populations, highlighting the deep genetic divergence between these oceanic regions (**Figure S5)**. However, NPA also showed evidence of shared ancestry with the Atlantic populations (ICE and SVA; **Figure S5)**. Increasing the number of clusters to K=4 identified four distinct genetic groups: one corresponding to the MED group, another to the SOC, a third encompassing the NPA population, and a fourth cluster that included both the ICE and SVA populations (Figure 2c-d). Interestingly, at K=5, the ICE and SVA populations remained distinct clusters from MED, although a few MED samples still showed shared ancestry with them (**Figure S5)**. Moreover, the NPA population exhibited signs of substructure within the Pacific Ocean basin

**Figure 2.**
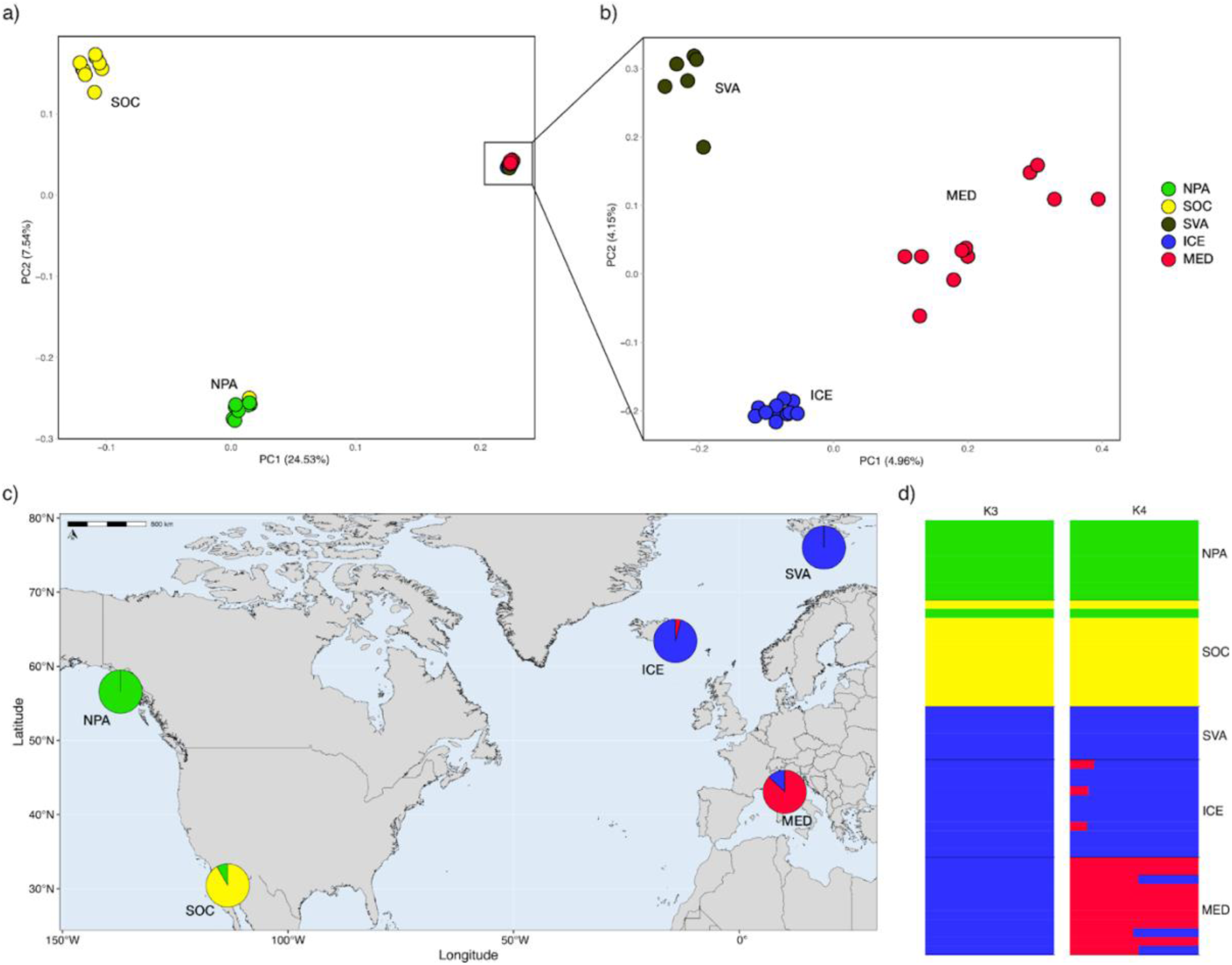
Population structure. PCA on the entire dataset (a) and excluding individuals from the Sea of Cortez (SOC) and the North Pacific (NPA) (b). (c, d) Admixture analyses. Each bar in panel d represents an individual, and each colour indicates the proportion of that individual’s genome assigned to each of the K clusters.

Pairwise F_ST_ analysis revealed a relatively high level of genetic differentiation between the SOC population and all other populations (above 15%) (**Table S9**). Moderate genetic differentiation was observed between the NPA population and the North Atlantic and Mediterranean populations (above 17%). In contrast, F_ST_ values among the MED, ICE, and SVA populations were consistently low, all falling below 3%.

### 3.4 Genomic diversity and inbreeding

Genome-wide diversity estimates revealed marked differences among fin whale populations. In particular, *θ*_W_ values ranged from highest in NPA to lowest in SOC (Figure 3a). Since ROH are genomic regions characterized by low variability, their inclusion in diversity estimates is expected to reduce genome-wide values. However, the extent of this reduction may differ across populations, reflecting variation in demographic history and levels of inbreeding. The inclusion of ROH regions led to a reduction in diversity estimates of 1.25% in NPA, 2.46% in SVA, 5.91% in ICE, 6.01% in MED, and a marked 35.58% in SOC, consistent with expectations for a small population subject to recent inbreeding (Figure 3a). Genomic diversity estimates in both neutral regions (outside CDS) and CDS showed a lower diversity in SOC and higher in NPA population (Figure 3b), while populations in the Atlantic Ocean and Mediterranean Sea showed similar values. The degree of divergence of *θ*_W_ estimate between neutral regions and CDS was higher in SVA (59.52%) and lower in MED (41.91%) (Figure 3b).

**Figure 3.**
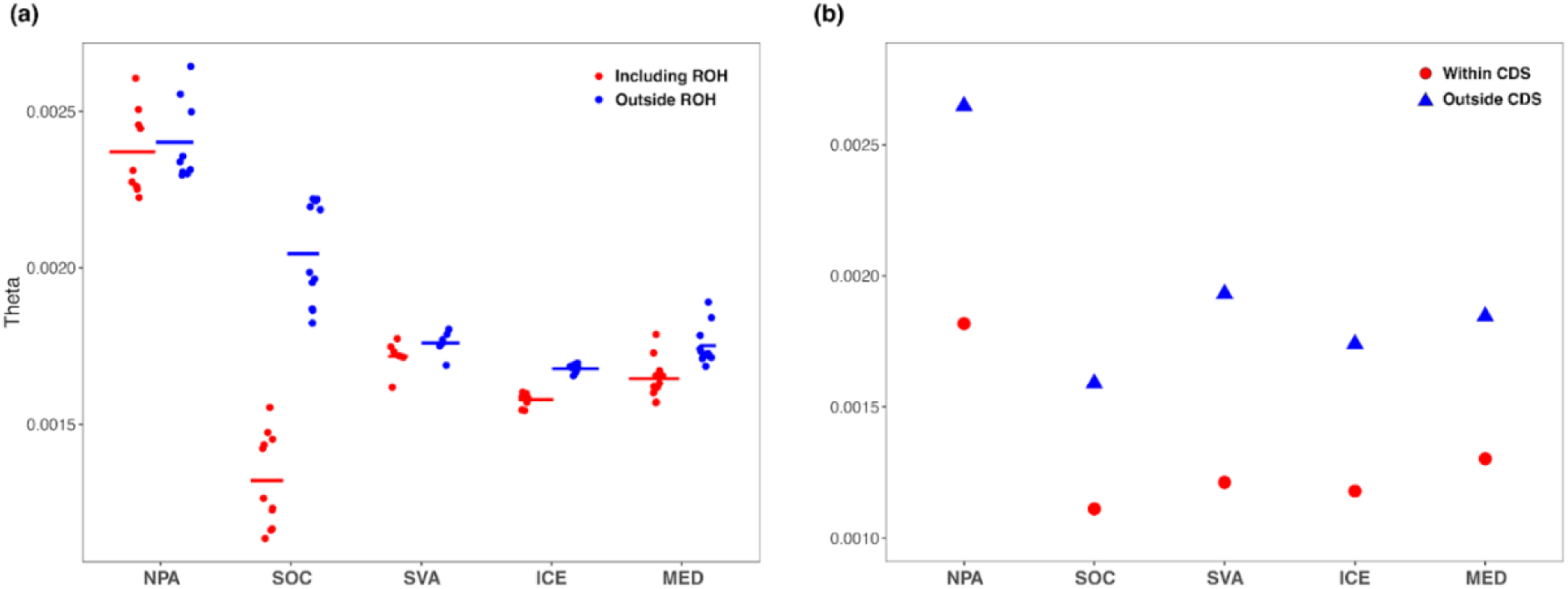
Genomic diversity estimates based on Watterson’s Theta *θ*_W_ including or excluding ROH regions (a) and estimates of *θ*_W_ per site within and outside exons (b).

The fraction of the genome being in ROH (FROH) was consistent within each population. The SOC population had the highest mean FROH (39.17%). Lower FROH levels were observed in the populations of MED, ICE and SVA, with a mean FROH value of 6.50%, 6.32% and 2.80%, respectively. The lowest level of FROH was registered for NPA that has a mean FROH value of 1.81% (Figure 4a). The individual SOC02, assigned to NPA by mitochondrial and nuclear analysis, has a mean FROH value comparable to other individuals belonging to the NPA population. In comparison to the other populations, the SOC population showed a high number of ROH larger than 2.5 Mb (Figure 4b). ICE and MED populations also showed a number of ROH above 2.5 Mb slightly higher than SVA and NPA (Figure 4b).

**Figure 4.**
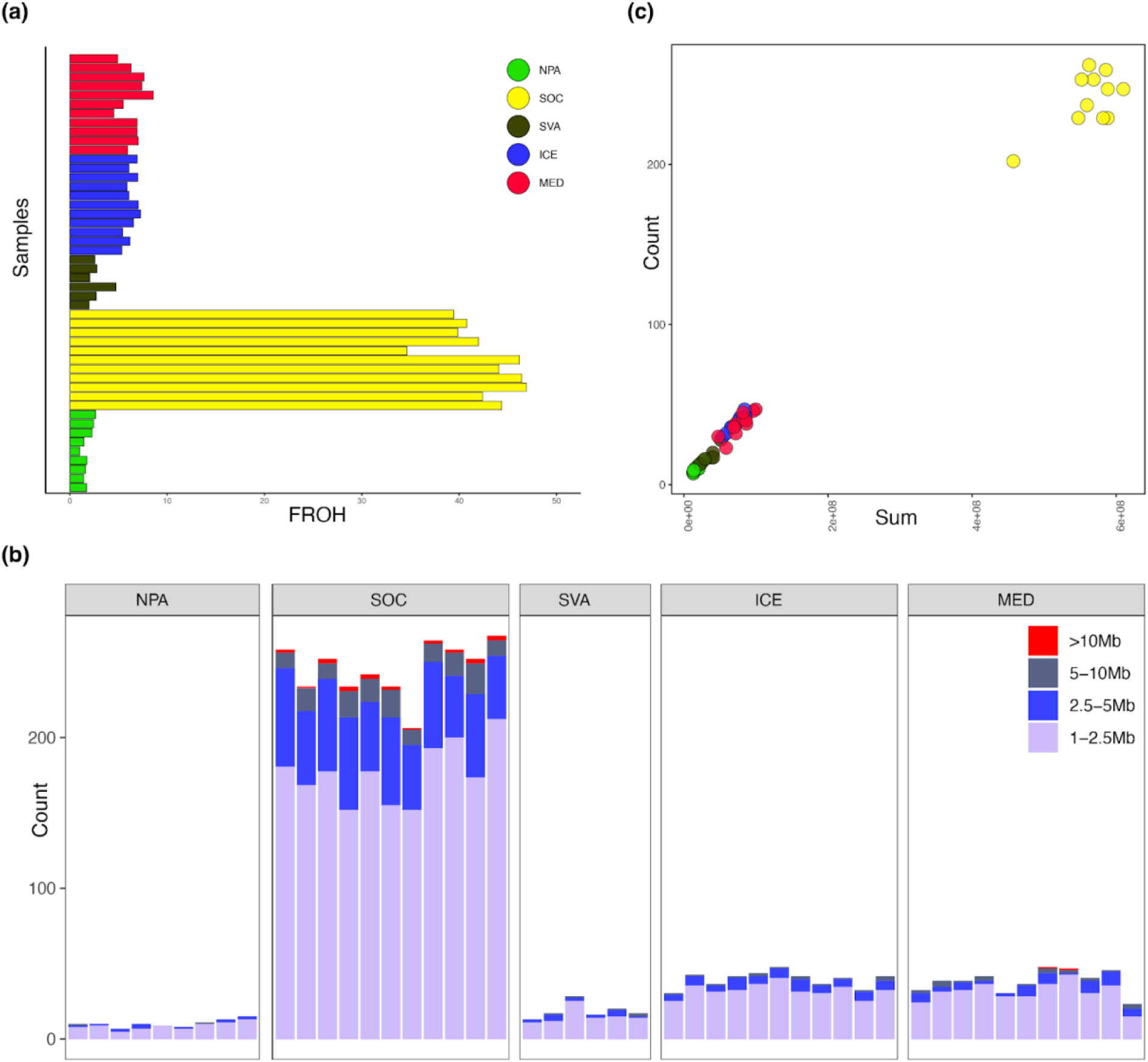
Inbreeding in fin whale populations. (b) The fraction of the genome in ROH (>1Mb) estimated with ROHan. Colour is representative of the ROH size in megabases (Mb). (c) Number of ROH compared to the sum of the length of ROH across the autosomes.

The sum and number of runs of homozygosity (SROH and NROH, respectively) are expected to be low in large populations, and both indices can be further reduced by admixture. In particular, admixture between highly divergent lineages carrying different haplotypes will reduce the number and sum of ROH in the genome (Ceballos, Joshi, et al., 2018). In contrast, populations that have experienced bottlenecks will have higher SROH and NROH (Ceballos, Joshi, et al., 2018). In our dataset SROH and NROH within individual genomes were highly correlated (Spearman correlation r = 0.879, p = 2.2e-16), with the SOC population exhibiting particularly high levels of both metrics. Among the MED population, some individuals from displayed relatively high values (Figure 4c). The SROH and NROH are expected to reflect population demography (Ceballos, Joshi, et al., 2018).

### 3.5 MHC *DQB-1* genetic variation

The variable regions of the MHC loci exemplify genomic regions in which greater intra- and inter-individual diversity reflects enhanced adaptation and response to pathogens (Bernachez and Landry 2003). Here we analysed exon 2, the most variable of the MHC *DQB-1* locus (Nigenda Morales et al. 2008), in the fin whale genomes from the different populations analysed in this work (NPA, SOC, SVA, ICE and MED), together with orthologous sequences from public repositories for two closely related species (*Balaenoptera musculus,* MUS and *Megaptera novaeangliae*, MEG) (**Table 2**). Basic indices of genetic diversity suggested that all analysed fin whale populations showed the expected level of variation at a key locus involved in pathogen interaction. In particular, in this genomic region, *θw*, a measure of the population mutation rate, was ten times higher than in the rest of the coding regions in the case of NPA, and even twenty times higher in all other cases (Figure 3). These results suggest that, similar to the other groups, a good level of adaptive MHC diversity is maintained in the MED population, as well as in the SOC population, which is known for its small effective population size (Ne).

**Table 2.** MHC *DQB-1* genetic diversity and results of Tajima’s *D* neutrality test.

|  | NPA | SOC | SVA | ICE | MED | MUS | MEG |
| --- | --- | --- | --- | --- | --- | --- | --- |
| sample size | 9 | 7 | 6 | 11 | 7 | 5 | 5 |
| variable sites | 27 | 26 | 28 | 30 | 31 | 13 | 14 |
| $\pi$ | 0.046 | 0.049 | 0.048 | 0.050 | 0.046 | 0.021 | 0.022 |
| $\theta_w$ | 0.029 | 0.030 | 0.034 | 0.030 | 0.036 | 0.017 | 0.018 |
| Tajima's *D* | 2.29* | 2.57** | 1.74 | 2.40* | 1.23 | 0.99 | 0.81 |
| *$p < 0.05$ **$p < 0.01$ | | | | | | | |

### 3.6 Demographic history

We modelled the effective population size (Ne) over the past 20 million years (Ma) using a multiple sequentially Markovian coalescent (MSMC) analysis (Schiffels and Durbin, 2014), based on the distribution of heterozygous sites across the genome (Figure 5). Ancestral effective population sizes, particularly for populations from the Pacific Ocean (NPA and SOC), were notably higher during the Plio-Pleistocene transition (LPT, 2.6 Ma ago) compared to recent estimates (Figure 5). After the mid-Pleistocene transition (MPT), Ne for all populations slightly increased until approximately 100-200 thousand years ago (ka), which coincides with the last interglacial periods. Following this time, fin whale populations in the Atlantic Ocean (ICE and SVA) and MED declined, whereas populations from NPA and SOC remained stable. More recently, the SOC population has experienced a decline.

**Figure 5.**
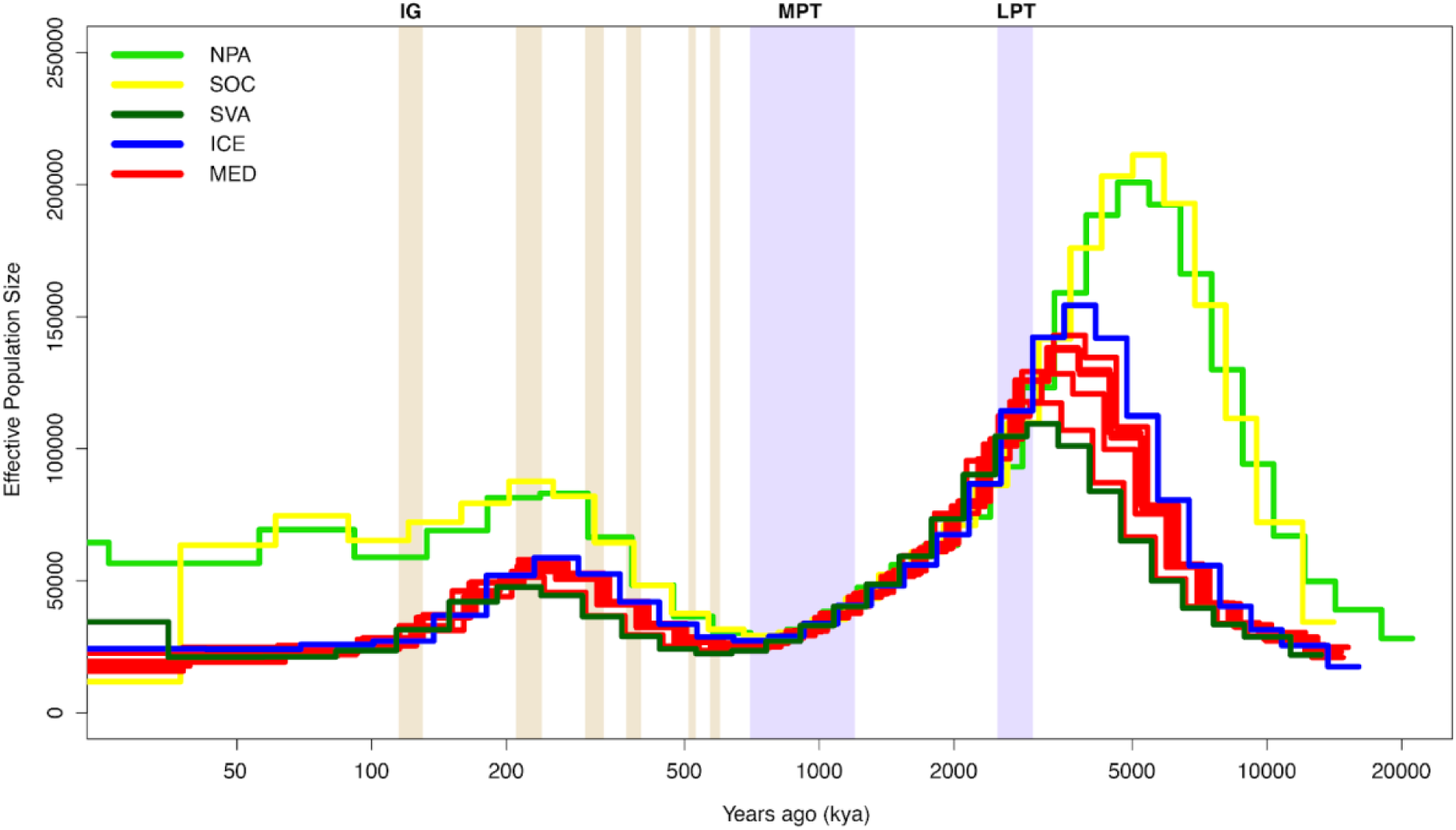
MSMC effective population size estimates. The x axis shows the time, and the y axis shows Ne. The model covers the last 20 Mya to 30 kya and is scaled based on a mutation rate of 1.38 × 10−8 per site per generation (REF) and a generation time of 25.9 years. The mid-Pleistocene transition (MPT, 0.7–1.2 million years ago, Ma) and the Late-Pleistocene Transition (LPT, 2.6 Ma) are shown as the light blue shaded region, light red shading indicates interglacials (IG) in the Pleistocene and Holocene.

### 3.7 Genetic load

We estimated the genetic load in all populations for samples with at least 10x coverage (N = 40; **Table S4-S5**) and used two outgroups (*B. musculus* and *M. novaeangliae*) to polarise each SNP as ancestral or derived. We first calculated the mutational load based on functional annotations from SNP effects predicted by snpEff (Cingolani et al., 2012). This allowed us to assess potentially deleterious variation in two impact categories: moderate impact (missense) and low impact (synonymous). Both categories exhibited a similar pattern of homozygous derived and heterozygous derived genotypes across the five populations (Figure 6a). For both mutation types, heterozygosity was reduced, while homozygosity was elevated in the SOC population. Additionally, we assessed the potentially deleterious effects of mutations using conservation scores derived from the alignment of homologous sequences from a large number of species. These scores provide a quantitative estimate of deleteriousness. When examining variants in coding regions, which are more likely to have functional consequences, we observed a similar trend based on GERP scores (Figure 6b). The MED population exhibited a lower masked load compared to NPA, but a higher realized load (Figure 6b). The higher realized load in SOC, reflecting an increase in homozygous deleterious alleles, suggested that these mutations are fully expressed in this population. This may be attributed to their smaller population size, where genetic drift likely plays a significant role. In smaller populations, genetic drift can lead to the fixation of deleterious alleles, thereby increasing the realized load (Bertorelle et al., 2022).

**Figure 6.**
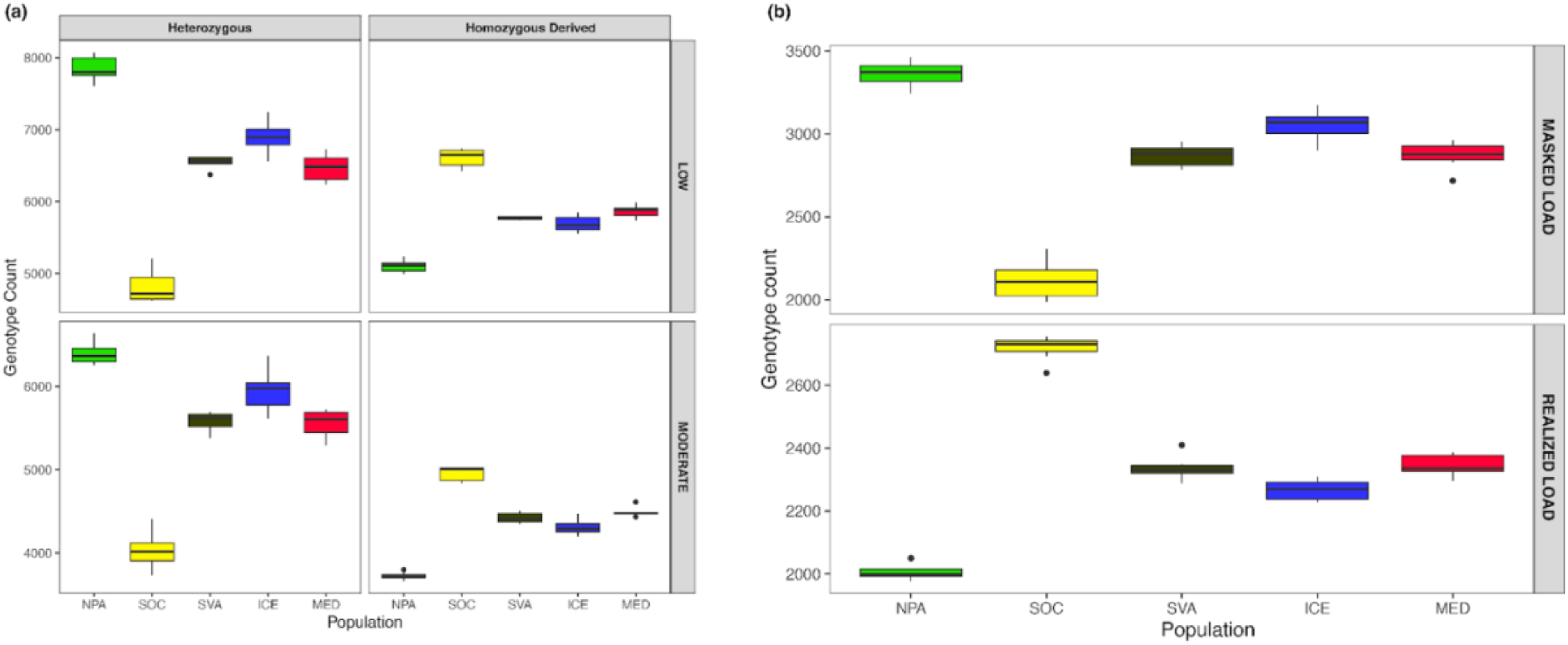
Genetic load. (a) Proportion of heterozygous genotypes and homozygous derived genotypes for synonymous sites (LOW) and missense sites (MODERATE) in different fin whale populations. (b) Genetic load divided into the components masked load (top) and realized load (bottom) estimated as the count of genotypes with GERP scores over all deleterious derived alleles in heterozygous and homozygous genotypes.

## 4 DISCUSSION

The Mediterranean fin whale population is classified as Endangered and has long been considered genetically distinct from the North Atlantic populations, although the degree of isolation and the boundaries between these populations remain subjects of debate.

Using a whole-genome approach, we found that Mediterranean fin whales are not completely genetically isolated. Although the Mediterranean population forms a distinct genetic cluster separated from the North Atlantic population, a more complex substructure emerged within the Mediterranean Sea. In particular, our results suggested that gene flow between the Mediterranean and North Atlantic populations persists, but it may preferentially involve a subgroup of individuals, with others being more isolated. This pattern supports occasional or seasonally migrations of fin whales through the Strait of Gibraltar suggested by previous studies (Gauffier et al., 2009, 2018; Bentaleb et al., 2011; Castellote et al., 2012). It could also agree with the hypothesis that some fin whales observed in the Mediterranean Sea may represent a summer feeding group that migrates elsewhere for winter breeding, potentially as part of the eastern North Atlantic population (e.g., in the Strait of Gibraltar). A certain degree of population structure within the Mediterranean Sea was also suggested by bioacoustics studies, which identified two contrasting song patterns of fin whales in the Mediterranean Sea, attributable to the presence of two different populations (Castellote et al. 2012; Sciacca et al. 2015; Pereira et al. 2020). Of these two populations, one has been described as migrating seasonally to the Atlantic through the Strait of Gibraltar, while the other as year-round resident in the Mediterranean Sea (Castellote et al. 2012; Di Sciara et al., 2016; Pereira et al. 2020). Our findings suggest a possible alignment with this division, hinting that the more genetically distinct individuals may represent a resident Mediterranean subpopulation (Figure 2).

However, the genetic divergence between North Atlantic (SVA and ICE) and MED fin whales was lower than that observed between other oceanic basins, such as the SOC and the NPA. This pattern aligns with a stepping-stone model of gene flow, where genetic exchange occurs principally between nearby regions, leading to increased differentiation over greater distances (Kimura and Weiss 1964). This is further supported by the lower genetic distance between ICE and both SVA and MED compared to the higher distance between SVA and MED (**Table S9**).

The MED population showed levels of heterozygosity and inbreeding comparable to those of the North Atlantic populations, in particular with ICE population, indicates that, although there may be some genetic isolation, the MED population is not yet facing extreme signs of genomic erosions, unlike the more isolated SOC population (Figure 3**-4**). One possible explanation is that, despite being limited, gene flow has been sufficient to maintain relatively stable genetic diversity. However, given the long generation times of the species and its recent decline, the MED population may be experiencing a drift debt, a time-lagged genetic effects caused by a recent population bottleneck or diminished gene flow. While the MED population is at present exhibiting similar levels of genetic diversity, the complete effects of restricted gene flow and historical demographic declines may not yet be entirely evident. Moreover, it is important to consider that the Eastern North Atlantic populations went through a severe genetic bottleneck due to extensive whaling which began thousands of years ago and peaked in the last two centuries (Aguilar and García-Vernet 2018; Wolf et al., 2021). The effects of this historical population decline may not yet be fully evident, and given the gene flow between the two basins, this could also influence the genetic diversity and inbreeding of the MED population that is already higher than other North Atlantic population (SVA).

Analysis of the MHC DQB-1 locus, a key genomic region involved in immune response, confirms that fin whales from the Mediterranean Sea maintain an adaptive level of genetic diversity comparable to other fin whale populations (**Table 2**). This suggests that, despite demographic challenges, they retain a relatively high genetic capacity for pathogen resistance, an important factor for long-term viability. However, the fate of this genetic diversity is difficult to predict if these populations become more isolated in the future and if the trend of decreasing population size continues. Although in some cases selection may counteract genetic drift and help maintain genetic variation at loci responsible for immunity against parasites (Robinson et al. 2016; Benazzo et al. 2017; Marmesat et al. 2017), several studies show that such diversity can be eroded during demographic decline (Sutton et al. 2011; Sutton et al. 2015; Zhang et al. 2016). In this context, the recent detection of morbillivirus infections in MED fin whales may have particularly severe consequences, since the virus induces immunosuppression also via MHC downregulation (Mazzariol et al 2016; Gaur et al. 2024). Low MHC variation, when combined with morbillivirus-induced downregulation of MHC expression, may critically impair antigen recognition and T cell activation, resulting in increased susceptibility to infection.

Our MSMC analysis revealed slightly different demographic histories between Pacific and Atlantic fin whale populations (Figure 5). Pacific populations (NPA and SOC) maintained larger ancestral effective population sizes (Ne), particularly during the Plio-Pleistocene transition (∼2.6 Ma). In contrast, Atlantic (ICE and SVA) and Mediterranean (MED) populations exhibited lower ancestral Ne and experienced a smaller recover following the mid-Pleistocene transition (∼700–200 ka). These different trends likely reflect differences in connectivity and environmental conditions across these ocean basins. Although MSMC is not accurate for recent time (Hilgers et al., 2025), the observed long-term decline in Ne for North Atlantic and MED populations highlights potential vulnerability to both past climatic shifts and present demographic constraints. These patterns agree with previous genomic studies (Yim et al., 2014; Arnason et al., 2018) and highlight the complex evolutionary histories of different fin whale populations.

The MED fin whales exhibited similar levels of masked and realized load with North Atlantic populations but differed from NPA and SOC populations (Figure 6). The MED population exhibited a lower masked load compared to NPA, but a higher realized load. This pattern has already been observed in the SOC and NPA populations of fin whales and is consistent with reduced genome-wide heterozygosity and small population sizes (Nigenda-Morales et al., 2023). What we report here for the first time is that populations from the North Atlantic (ICE and SVA) and MED showed similar levels among themselves and an intermediate genetic load, between SOC (small population) and NPA (large population). The higher realized load in SOC, reflecting an increase in homozygous deleterious alleles, suggested that these mutations are fully expressed in this population. These results implies that SOC individuals have genomes carrying a higher realized load compared to MED population. However, future consanguineous mating would unmask more deleterious mutations in MED and that their relatively low effective population size and restricted connectivity put them at risk if environmental or demographic pressures intensify.

Despite some ongoing gene flow with the North Atlantic and comparable level of genomic diversity, inbreeding, genetic load and adaptive variation, the Mediterranean fin whale population faces substantial risks due to its small size, partial isolation, and vulnerability to environmental changes. Rising temperatures and anthropogenic stressors may disrupt migration, feeding, and reproduction, leading to a lower input of novel genetic variants, affecting both potential evolutionary change and genetic load (Kebke et al., 2022; Silber et al., 2017; Nigenda-Morales et al., 2023; Bertorelle et al., 2022). Moreover, if resident individuals will be forced to alter their geographic range, as observed in other fin whale populations (Ruiz-Sagales et al., 2024), behavioural plasticity, including shifts in diet and migration, may enable Mediterranean fin whales to adapt to new habitats potentially causing an imbalance in the ecosystem they currently belong to (Pearson et al., 2023).

Conservation efforts should prioritize protecting Mediterranean fin whale habitats, reducing human-induced threats such as ship strikes, noise pollution, and climate-driven shifts in prey availability. Future studies should continue to monitor genetic diversity and migration patterns by expanding the area of sampling, as well as assess how environmental changes impact the delicate balance between Mediterranean and North Atlantic fin whale populations.

## ACKNOWLEDGEMENTS

Support was provided to AI by the Italian Ministry of University and Research through the National Biodiversity Future Center, part of the National Recovery and Resilience Plan, Mission 4, Component 2, Investment 1.4, Project CN00000033. This study was also funded to MCF by the Interreg MED project: Plastic Busters MPAs: Preserving Biodiversity from Plastics in Mediterranean Marine Protected Areas, co-funded by the European Regional Development Fund (grant agreement No 4MED17_3.2_M123_027). Samples from Mexico were collected under the permits SGPA/DGVS/000612/18 and SGPA/DGVS/013212/18. A special thanks to CIMA Research Foundation and ISPRA for their contribution in fin whale sampling in the Mediterranean Sea. We thank also Alessia Profico for helping with the genome annotation.

## DATA AVAILABILITY

The genome assembly and the Illumina reads of samples sequenced in this study were deposited in the National Center for Biotechnology Information (NCBI) with BioProject number PRJNA1268741. Genome assembly and annotation files are deposited on Zenodo (10.5281/zenodo.15649665). Scripts are deposited on GitHub (https://github.com/rsbiello/FinWhale_popgen).

## AUTHOR CONTRIBUTIONS

RB, SF, CP, and AM conceived and designed the study. JUR, MCF and CF provided samples. AI, ED and CP carried out laboratory work. RB, AI, SF and ED analysed the data. RB, AI, SF and ED wrote the paper with input from all other co-authors.

## REFERENCES

Aguilar, A., & García-Vernet, R. (2018). Fin whale: *Balaenoptera physalus*. In Encyclopedia of marine mammals (pp. 368-371). Academic Press.

Andrews, S. (2010). FastQC: a quality control tool for high throughput sequence data. Available online at: http://www.bioinformatics.babraham.ac.uk/projects/fastqc.

Archer, F. I., Morin, P. A., Hancock-Hanser, B. L., Robertson, K. M., Leslie, M. S., Bérubé, M., … & Taylor, B. L. (2013). Mitogenomic phylogenetics of fin whales (*Balaenoptera physalus* spp.): genetic evidence for revision of subspecies. PLoS One, 8(5), e63396.

Árnason, U. (1969). The karyotype of the fin whale. Hereditas, 62(3), 273–284.

Árnason, Ú., Spilliaert, R., Pálsdóttir, Á., & Árnason, A. (1991). Molecular identification of hybrids between the two largest whale species, the blue whale (*Balaenoptera musculus*) and the fin whale (*B. physalus*). Hereditas, 115(2), 183–189.

Árnason, Ú., Lammers, F., Kumar, V., Nilsson, M. A., & Janke, A. (2018). Whole-genome sequencing of the blue whale and other rorquals finds signatures for introgressive gene flow. Science advances, 4(4), eaap9873.

Benazzo, A., Trucchi, E., Cahill, J. A., Maisano Delser, P., Mona, S., Fumagalli, M., … & Bertorelle, G. (2017). Survival and divergence in a small group: The extraordinary genomic history of the endangered Apennine brown bear stragglers. Proceedings of the National Academy of Sciences, 114(45), E9589–E9597.

Bentaleb, I., Martin, C., Vrac, M., Mate, B., Mayzaud, P., Siret, D., … & Guinet, C. (2011). Foraging ecology of Mediterranean fin whales in a changing environment elucidated by satellite tracking and baleen plate stable isotopes. Marine Ecology Progress Series, 438, 285–302.

Bernatchez, L., & Landry, C. (2003). MHC studies in nonmodel vertebrates: what have we learned about natural selection in 15 years? Journal of evolutionary biology, 16(3), 363–377.

Bertorelle, G., Raffini, F., Bosse, M., Bortoluzzi, C., Iannucci, A., Trucchi, E., … & Van Oosterhout, C. (2022). Genetic load: genomic estimates and applications in non-model animals. Nature Reviews Genetics, 23(8), 492–503.

Bérubé, M., Aguilar, A., Dendanto, D., Larsen, F., Notarbartolo Di Sciara, G., Sears, R., … & Palsbøll, P. J. (1998). Population genetic structure of North Atlantic, Mediterranean Sea and Sea of Cortez fin whales, Balaenoptera physalus (Linnaeus 1758): analysis of mitochondrial and nuclear loci. Molecular ecology, 7(5), 585–599.

Bérubé, M., Urbán, J., Dizon, A. E., Brownell, R. L., & Palsbøll, P. J. (2002). Genetic identification of a small and highly isolated population of fin whales (*Balaenoptera physalus*) in the Sea of Cortez, Mexico. Conservation Genetics, 3, 183–190.

Bhatia, G., Patterson, N., Sankararaman, S., & Price, A. L. (2013). Estimating and interpreting FST: The impact of rare variants. Genome Research, 23(9), 1514–1521.

Bryant, D., & Moulton, V. (2004). Neighbor-net: an agglomerative method for the construction of phylogenetic networks. Molecular Biology and Evolution, 21(2), 255–265. 10.1093/molbev/msh018

Brůna, T., Hoff, K. J., Lomsadze, A., Stanke, M., & Borodovsky, M. (2021). BRAKER2: automatic eukaryotic genome annotation with GeneMark-EP+ and AUGUSTUS supported by a protein database. NAR genomics and bioinformatics, 3(1), lqaa108.

Buchfink, B., Xie, C., & Huson, D. H. (2015). Fast and sensitive protein alignment using DIAMOND. Nature methods, 12(1), 59–60.

Buss, D. L., Atmore, L. M., Zicos, M. H., Goodall-Copestake, W. P., Brace, S., Archer, F. I., … & Jackson, J. A. (2023). Historical Mitogenomic Diversity and Population Structuring of Southern Hemisphere Fin Whales. Genes, 14(5), 1038.

Cabrera, A. A., Bérubé, M., Lopes, X. M., Louis, M., Oosting, T., Rey-Iglesia, A., … & Palsbøll, P. J. (2021). A genetic perspective on cetacean evolution. Annual Review of Ecology, Evolution, and Systematics, 52, 131–151.

Cabrera, A. A., Hoekendijk, J. P., Aguilar, A., Barco, S. G., Berrow, S., Bloch, D., … & Bérubé, M. (2019). Fin whale (*Balaenoptera physalus*) mitogenomics: A cautionary tale of defining sub-species from mitochondrial sequence monophyly. Molecular phylogenetics and evolution, 135, 86–97.

Castellote, M., Clark, C. W., & Lammers, M. O. (2012). Fin whale (*Balaenoptera physalus*) population identity in the western Mediterranean Sea. Marine Mammal Science, 28(2), 325–344.

Ceballos, F. C., Joshi, P. K., Clark, D. W., Ramsay, M., & Wilson, J. F. (2018). Runs of homozygosity: windows into population history and trait architecture. Nature Reviews Genetics, 19(4), 220–234.

Cingolani, P., Platts, A., Wang, L. L., Coon, M., Nguyen, T., Wang, L., Land, S., Lu, X., & Ruden, D. M. (2012). A program for annotating and predicting the effects of single nucleotide polymorphisms, SnpEff: SNPs in the genome of Drosophila melanogaster strain w1118; iso-2; iso-3. fly, 6(2), 80–92.

Clapham, P. J., Young, S. B., & Brownell Jr, R. L. (1999). Baleen whales: conservation issues and the status of the most endangered populations. Mammal review, 29(1), 37–62.

Coll, M., Piroddi, C., Albouy, C., Ben Rais Lasram, F., Cheung, W. W., Christensen, V., … & Pauly, D. (2012). The Mediterranean Sea under siege: spatial overlap between marine biodiversity, cumulative threats and marine reserves. Global Ecology and Biogeography, 21(4), 465–480.

Collignon, A., Hecq, J. H., Glagani, F., Voisin, P., Collard, F., & Goffart, A. (2012). Neustonic microplastic and zooplankton in the North Western Mediterranean Sea. Marine pollution bulletin, 64(4), 861–864.

Conesa, A., Götz, S., García-Gómez, J. M., Terol, J., Talón, M., and Robles, M. (2005). Blast2GO: a universal tool for annotation, visualization and analysis in functional genomics research. Bioinformatics, 21, 3674–3676. doi: 10.1093/bioinformatics/bti610

Cotté, C., d’Ovidio, F., Chaigneau, A., Lévy, M., Taupier-Letage, I., Mate, B., & Guinet, C. (2011). Scale-dependent interactions of Mediterranean whales with marine dynamics. Limnology and Oceanography, 56(1), 219–232.

Cózar, A., Sanz-Martín, M., Martí, E., González-Gordillo, J. I., Ubeda, B., Gálvez, J. Á., … & Duarte, C. M. (2015). Plastic accumulation in the Mediterranean Sea. PloS One, 10(4), e0121762.

Das, K., Holleville, O., Ryan, C., Berrow, S., Gilles, A., Ody, D., & Michel, L. N. (2017). Isotopic niches of fin whales from the Mediterranean Sea and the Celtic Sea (North Atlantic). Marine Environmental Research, 127, 75–83.

Danecek, P., Auton, A., Abecasis, G., Albers, C.A., Banks, E., DePristo, M.A. et al. (2011). The variant call format and VCFtools. Bioinformatics, 27, 2156–2158.

Danecek, P., Bonfield, J.K., Liddle, J., Marshall, J., Ohan, V., Pollard, M.O. et al. (2021). Twelve years of SAMtools and BCFtools. Gigascience, 10, giab008.

Derrien, T., Estellé, J., Marco Sola, S., Knowles, D. G., Raineri, E., Guigó, R., & Ribeca, P. (2012). Fast computation and applications of genome mappability. PloS One, 7(1), e30377.

Di Sciara, G. N., Castellote, M., Druon, J. N., & Panigada, S. (2016). Fin whales, *Balaenoptera physalus*: at home in a changing Mediterranean Sea? Advances in Marine Biology, 75, 75–101.

Di Sciara, G. N., Venturino, M. C., Zanardelli, M., Bearzi, G., Borsani, F. J., & Cavalloni, B. (1993). Cetaceans in the central Mediterranean Sea: distribution and sighting frequencies. Italian Journal of Zoology, 60(1), 131–138.

Di Sciara, G. N., Zanardelli, M., Jahoda, M., Panigada, S., & Airoldi, S. (2003). The fin whale Balaenoptera physalus (L. 1758) in the Mediterranean Sea. Mammal Review, 33(2), 105–150.

Dudchenko, O., Shamim, M. S., Batra, S. S., Durand, N. C., Musial, N. T., Mostofa, R., … & Aiden, E. L. (2018). The Juicebox Assembly Tools module facilitates de novo assembly of mammalian genomes with chromosome-length scaffolds for under $1000. BioRxiv, 254797.

Durfort, A., Mariani, G., Tulloch, V., Savoca, M. S., Troussellier, M., & Mouillot, D. (2022). Recovery of carbon benefits by overharvested baleen whale populations is threatened by climate change. Proceedings of the Royal Society B, 289(1986), 20220375.

Edwards, E. F., Hall, C., Moore, T. J., Sheredy, C., & Redfern, J. V. (2015). Global distribution of fin whales *Balaenoptera physalus* in the post-whaling era (1980–2012). Mammal Review, 45(4), 197–214.

Emms, D. M., & Kelly, S. (2019). OrthoFinder: phylogenetic orthology inference for comparative genomics. Genome Biology, 20, 1–14.

Espada, R., Camacho-Sánchez, A., Olaya-Ponzone, L., Martín-Moreno, E., Patón, D., & García-Gómez, J. C. (2024). Fin Whale Balaenoptera physalus Historical Sightings and Strandings, Ship Strikes, Breeding Areas and Other Threats in the Mediterranean Sea: A Review (1624–2023). Environments, 11(6), 104.

Fan, G., Zhang, Y., Liu, X., Wang, J., Sun, Z., Sun, S., Zhang, H. E., Chen, J., Lv, M., Han, K., Tan, X., Hu, J., Guan, R., Fu, Y., Liu, S., Chen, X. I., Xu, Q., Qin, Y., Liu, L., … Liu, X. (2019). The first chromosome-level genome for a marine mammal as a resource to study ecology and evolution. Molecular Ecology Resources, 19(4), 944–956.

Finn RD, Bateman A, Clements J, Coggill P, Eberhardt RY, Eddy SR, Heger A, Hetherington K, Holm L, Mistry J (2013) Pfam: the protein families database. Nucleic Acids Res, 42, D222–D230.

Forney, K. A., & Barlow, J. (1998). Seasonal patterns in the abundance and distribution of California cetaceans, 1991–1992. Marine Mammal Science, 14(3), 460–489.

Fossi, M. C., Baini, M., & Simmonds, M. P. (2020). Cetaceans as ocean health indicators of marine litter impact at global scale. Frontiers in Environmental Science, 8, 586627.

Fossi, M. C., Marsili, L., Baini, M., Giannetti, M., Coppola, D., Guerranti, C., … & Panti, C. (2016). Fin whales and microplastics: The Mediterranean Sea and the Sea of Cortez scenarios. Environmental Pollution, 209, 68–78.

Fossi, M. C., Panti, C., Marsili, L., Maltese, S., Spinsanti, G., Casini, S., … & Finoia, M. G. (2013). The Pelagos Sanctuary for Mediterranean marine mammals: Marine Protected Area (MPA) or marine polluted area? The case study of the striped dolphin (*Stenella coeruleoalba*). Marine pollution bulletin, 70(1-2), 64–72.

Gabriel, L., Hoff, K. J., Bruna, T., Lomsadze, A., Borodovsky, M., & Stanke, M. (2023, January). The BRAKER3 genome annotation pipeline. In Plant and Animal Genomes Conference (Vol. 30).

Gabriel, L., Hoff, K. J., Brůna, T., Borodovsky, M., & Stanke, M. (2021). TSEBRA: transcript selector for BRAKER. BMC bioinformatics, 22, 1–12.

Ganley, L. C., Byrnes, J., Pendleton, D. E., Mayo, C. A., Friedland, K. D., Redfern, J. V., … & Brault, S. (2022). Effects of changing temperature phenology on the abundance of a critically endangered baleen whale. Global Ecology and Conservation, 38, e02193.

Garcia-Erill, G., & Albrechtsen, A. (2020). Evaluation of model fit of inferred admixture proportions. Molecular Ecology Resources, 20(4), 936–949.

Gauffier, P., Verborgh, P., Andreu, E., Esteban, R., Medina, B., Gallego, P., & De Stephanis, R. (2009). An update on fin whales (Balaenoptera physalus) migration through intense maritime traffic in the Strait of Gibraltar. Int Whaling Comm SC/61/BC6, Madeira.

Gauffier, P., Verborgh, P., Giménez, J., Esteban, R., Sierra, J. M. S., & de Stephanis, R. (2018). Contemporary migration of fin whales through the Strait of Gibraltar. Marine Ecology Progress Series, 588, 215–228.

Gaur, S. K., Jain, J., Chaudhary, Y., & Kaul, R. (2024). Insights into the mechanism of Morbillivirus induced immune suppression. Virology, 110212.

Hanghøj, K., Moltke, I., Andersen, P. A., Manica, A., & Korneliussen, T. S. (2019). Fast and accurate relatedness estimation from high-throughput sequencing data in the presence of inbreeding. GigaScience, 8(5), giz034.

Hedrick, P. W., & Lacy, R. C. (2015). Measuring relatedness between inbred individuals. Journal of Heredity, 106(1), 20–25.

Hilgers, L., Liu, S., Jensen, A., Brown, T., Cousins, T., Schweiger, R., … & Hiller, M. (2025). Avoidable false PSMC population size peaks occur across numerous studies. Current Biology, 35(4), 927–930.

Hoff, K. J., Lange, S., Lomsadze, A., Borodovsky, M., & Stanke, M. (2016). BRAKER1: unsupervised RNA-Seq-based genome annotation with GeneMark-ET and AUGUSTUS. Bioinformatics, 32(5), 767–769.

Hoff, K. J., Lomsadze, A., Borodovsky, M., & Stanke, M. (2019). Whole-genome annotation with BRAKER. Gene prediction: methods and protocols, 65–95.

Huson, D. H., & Bryant, D. (2006). Application of phylogenetic networks in evolutionary studies. Molecular Biology and Evolution, 23(2), 254–267.

Jurka, J., Kapitonov, V. V., Pavlicek, A., Klonowski, P., Kohany, O., & Walichiewicz, J. (2005). Repbase Update, a database of eukaryotic repetitive elements. Cytogenetic and Genome Research, 110(1-4), 462–467.

Katoh, K., & Standley, D. M. (2013). MAFFT multiple sequence alignment software version 7: improvements in performance and usability. Molecular Biology and Evolution, 30(4), 772–780.

Katona, S. K., & Whitehead, H. P. (1981). Identifying humpback whales using their natural markings. Polar Record, 20(128), 439–444.

Kearse, M., Moir, R., Wilson, A., Stones-Havas, S., Cheung, M., Sturrock, S., Buxton, S., Cooper, A., Markowitz, S., Duran, C., Thierer, T., Ashton, B., Meintjes, P., & Drummond, A. (2012). Geneious Basic: An integrated and extendable desktop software platform for the organization and analysis of sequence data. Bioinformatics, 28(12), 1647–1649.

Kebke, A., Samarra, F., & Derous, D. (2022). Climate change and cetacean health: impacts and future directions. Philosophical Transactions of the Royal Society B, 377(1854), 20210249.

Kellogg, R. (1929) What is known of the migrations of some of the whalebone whales? Smithsonian Institution. Annual Report of the Board of Regents, 467–494.

Kim, D., Langmead, B., & Salzberg, S. L. (2015). HISAT: a fast spliced aligner with low memory requirements. Nature Methods, 12(4), 357–360.

Kimura, M., & Weiss, G. H. (1964). The stepping stone model of population structure and the decrease of genetic correlation with distance. Genetics, 49(4), 561.

Korneliussen, T.S., Albrechtsen, A., & Nielsen, R. (2014). ANGSD: analysis of next generation sequencing data. BMC bioinformatics, 15, 1–3.

Li, H., & Durbin, R. (2009). Fast and accurate short read alignment with Burrows–Wheeler transform. bioinformatics, 25(14), 1754–1760.

Li, H., Handsaker, B., Wysoker, A., Fennell, T., Ruan, J., Homer, N., Marth G., Abecasis G., Durbin R., & 1000 Genome Project Data Processing Subgroup. (2009). The sequence alignment/map format and SAMtools. bioinformatics, 25(16), 2078–2079.

Marmesat, E., Schmidt, K., Saveljev, A. P., Seryodkin, I. V., & Godoy, J. A. (2017). Retention of functional variation despite extreme genomic erosion: MHC allelic repertoires in the *Lynx* genus. BMC Evolutionary Biology, 17, 1–16.

Mazzariol, S., Centelleghe, C., Beffagna, G., Povinelli, M., Terracciano, G., Cocumelli, C., … & Di Guardo, G. (2016). Mediterranean fin whales (*Balaenoptera physalus*) threatened by dolphin morbillivirus. Emerging infectious diseases, 22(2), 302.

McKenna, A., Hanna, M., Banks, E., Sivachenko, A., Cibulskis, K., Kernytsky, A., … DePristo, M. A. (2010). The Genome Analysis Toolkit: a MapReduce framework for analyzing next-generation DNA sequencing data. Genome Research, 20(9), 1297–1303. 10.1101/gr.107524.110

Meisner, J., & Albrechtsen, A. (2018). Inferring population structure and admixture proportions in low-depth NGS data. Genetics, 210(2), 719–731. 10.1534/genetics.118.301336

Mizroch, S. A., Rice, D. W., Zwiefelhofer, D., Waite, J., & Perryman, W. L. (2009). Distribution and movements of fin whales in the North Pacific Ocean. Mammal Review, 39(3), 193–227.

Moore, S. E., Stafford, K. M., Mellinger, D. K., & Hildebrand, J. A. (2006). Listening for large whales in the offshore waters of Alaska. BioScience, 56(1), 49–55.

Nigenda-Morales S, Flores-Ramírez S, Urbán-R J, Vázquez-Juárez R. MHC DQB-1 polymorphism in the Gulf of California fin whale (*Balaenoptera physalus*) population. Journal of Heredity, 99(1), 14–21. doi: 10.1093/jhered/esm087

Nigenda-Morales, S. F., Lin, M., Nuñez-Valencia, P. G., Kyriazis, C. C., Beichman, A. C., Robinson, J. A., … & Wayne, R. K. (2023). The genomic footprint of whaling and isolation in fin whale populations. Nature Communications, 14(1), 5465.

Pallin, L. J., Kellar, N. M., Steel, D., Botero-Acosta, N., Baker, C. S., Conroy, J. A., … & Friedlaender, A. S. (2023). A surplus no more? Variation in krill availability impacts reproductive rates of Antarctic baleen whales. Global change biology, 29(8), 2108–2121.

Palsbøll, P. J., Bérubé, M., Aguilar, A., Notarbartolo-Di-Sciara, G., & Nielsen, R. (2004). Discerning between recurrent gene flow and recent divergence under a finite-site mutation model applied to North Atlantic and Mediterranean Sea fin whale (*Balaenoptera physalus*) populations. Evolution, 58(3), 670–675.

Panigada, S., Donovan, G. P., Druon, J. N., Lauriano, G., Pierantonio, N., Pirotta, E., … & di Sciara, G. N. (2017). Satellite tagging of Mediterranean fin whales: working towards the identification of critical habitats and the focussing of mitigation measures. Scientific Reports, 7(1), 3365.

Panigada, S., Gauffier, P., & Notarbartolo di Sciara, G. (2021). Balaenoptera physalus (Mediterranean subpopulation). The IUCN red list of threatened species 2021: e. T16208224A50387979.

Panigada, S., Lauriano, G., Burt, L., Pierantonio, N., & Donovan, G. (2011). Monitoring winter and summer abundance of cetaceans in the Pelagos Sanctuary (northwestern Mediterranean Sea) through aerial surveys. PloS One, 6(7), e22878.

Panti, C., Baini, M., Lusher, A., Hernandez-Milan, G., Rebolledo, E. L. B., Unger, B., … & Fossi, M. C. (2019). Marine litter: One of the major threats for marine mammals. Outcomes from the European Cetacean Society workshop. Environmental pollution, 247, 72–79.

Pearson, H. C., Savoca, M. S., Costa, D. P., Lomas, M. W., Molina, R., Pershing, A. J., … & Roman, J. (2023). Whales in the carbon cycle: can recovery remove carbon dioxide? Trends in Ecology & Evolution, 38(3), 238–249.

Pereira, A., Harris, D., Tyack, P., & Matias, L. (2020). Fin whale acoustic presence and song characteristics in seas to the southwest of Portugal. The Journal of the Acoustical Society of America, 147(4), 2235–2249.

Pertea, M., Pertea, G. M., Antonescu, C. M., Chang, T. C., Mendell, J. T., & Salzberg, S. L. (2015). StringTie enables improved reconstruction of a transcriptome from RNA-seq reads. Nature biotechnology, 33(3), 290–295.

Peters, K. J., Stockin, K. A., & Saltré, F. (2022). On the rise: Climate change in New Zealand will cause sperm and blue whales to seek higher latitudes. Ecological Indicators, 142, 109235.

Punta M, Coggill PC, Eberhardt RY, Mistry J, Tate J, Boursnell C, Pang N, Forslund K, Ceric G, Clements J, Heger A, Holm L, Sonnhammer ELL, Eddy SR, Bateman A, Finn RD (2012) The Pfam protein families database. Nucleic acids research, 40(D1), D290–D301.

Quevillon, E., Silventoinen, V., Pillai, S., Harte, N., Mulder, N., Apweiler, R., & Lopez, R. (2005). InterProScan: protein domains identifier. Nucleic acids research, 33(suppl_2), W116-W120.

Quinlan, A. R., & Hall, I. M. (2010). BEDTools: a flexible suite of utilities for comparing genomic features. Bioinformatics, 26(6), 841–842.

Renaud, G., Hanghøj, K., Korneliussen, T. S., Willerslev, E., & Orlando, L. (2019). Joint estimates of heterozygosity and runs of homozygosity for modern and ancient samples. Genetics, 212(3), 587–614.

Robinson, J. A., Ortega-Del Vecchyo, D., Fan, Z., Kim, B. Y., Marsden, C. D., Lohmueller, K. E., & Wayne, R. K. (2016). Genomic flatlining in the endangered island fox. Current Biology, 26(9), 1183–1189.

Ruiz-Sagalés, M., García-Vernet, R., Sanchez-Espigares, J., Halldórsson, S. D., Chosson, V., Sigurðsson, G. M., … & Aguilar, A. (2024). Baleen stable isotopes reveal climate-driven behavioural shifts in North Atlantic fin whales. Science of the Total Environment, 955, 177164.

Schiffels, S., & Durbin, R. (2014). Inferring human population size and separation history from multiple genome sequences. Nature genetics, 46(8), 919–925.

Schiffels, S., & Wang, K. (2020). MSMC and MSMC2: the multiple sequentially Markovian coalescent. In Statistical population genomics (pp. 147-165). Humana.

Schliep, K. P. (2011). phangorn: phylogenetic analysis in R. Bioinformatics, 27(4), 592–593. 10.1093/bioinformatics/btq706

Schubert, M., Lindgreen, S., & Orlando, L. (2016). AdapterRemoval v2: rapid adapter trimming, identification, and read merging. BMC Research Notes, 9(1), 1–7. 10.1186/s13104-016-1900-2

Sciacca, V., Caruso, F., Beranzoli, L., Chierici, F., De Domenico, E., Embriaco, D., … & Riccobene, G. (2015). Annual acoustic presence of fin whale (*Balaenoptera physalus*) offshore eastern Sicily, central Mediterranean Sea. PloS one, 10(11), e0141838.

Silber, G. K., Lettrich, M. D., Thomas, P. O., Baker, J. D., Baumgartner, M., Becker, E. A., … & Waples, R. S. (2017). Projecting marine mammal distribution in a changing climate. Frontiers in Marine Science, 4, 413.

Simão, F. A., Waterhouse, R. M., Ioannidis, P., Kriventseva, E. V., & Zdobnov, E. M. (2015). BUSCO: assessing genome assembly and annotation completeness with single-copy orthologs. Bioinformatics, 31(19), 3210–3212.

Simmonds, M. P., & Eliott, W. J. (2009). Climate change and cetaceans: concerns and recent developments. Journal of the Marine biological Association of the United Kingdom, 89(1), 203–210.

Skotte, L., Korneliussen, T. S., & Albrechtsen, A. (2013). Estimating individual admixture proportions from next generation sequencing data. Genetics, 195(3), 693–702.

Smit, A., Hubley, R., & Green, P. (2013). RepeatMasker Open-4.0. http://www.repeatmasker.org

Stafford, K.M., Mellinger, D.K., Moore, S.E., & Fox, C.G. (2007). Seasonal variability and detection range modeling of baleen whale calls in the Gulf of Alaska, 1999–2002. Journal of the Acoustical Society of America, 122, 3378–3390.

Stamatakis, A. (2014). RAxML version 8: a tool for phylogenetic analysis and post-analysis of large phylogenies. Bioinformatics, 30(9), 1312–1313.

Sutton, J. T., Nakagawa, S., Robertson, B. C., & Jamieson, I. G. (2011). Disentangling the roles of natural selection and genetic drift in shaping variation at MHC immunity genes. Molecular Ecology, 20(21), 4408–4420.

Sutton, J. T., Robertson, B. C., & Jamieson, I. G. (2015). MHC variation reflects the bottleneck histories of New Zealand passerines. Molecular Ecology, 24(2), 362–373.

Tershy, B.R., Urbán, R.J., Breese, D., Rojas, B. L., Findley LY (1993) Are fin whales resident to the Gulf of California? Revista de Investigación Científica, 1, 69–72

Tollis, M., Robbins, J., Webb, A. E., Kuderna, L. F. K., Caulin, A. F., Garcia, J. D., Bèrubè, M., Pourmand, N., Marques-Bonet, T., O’Connell, M. J., Palsbøll, P. J., & Maley, C. C. (2019). Return to the sea, get huge, beat cancer: An analysis of cetacean genomes including an assembly for the humpback whale (*Megaptera novaeangliae*). Molecular Biology and Evolution, 36(8), 1746–1763. 10.1093/molbev/msz099

Tulloch, V. J., Plagányi, É. E., Brown, C., Richardson, A. J., & Matear, R. (2019). Future recovery of baleen whales is imperiled by climate change. Global change biology, 25(4), 1263–1281.

Urbán, R. J., Rojas-Bracho, L., Guerrero-Ruiz, M., Jaramillo-Legorreta, A. & Findley, L. T. (2005). Cetacean diversity and conservation in the Gulf of California. Biodiversity, ecosystems, and conservation in Northern Mexico, 276–297.

Watterson, G. A. (1975). On the number of segregating sites in genetical models without recombination. Theoretical population biology, 7(2), 256–276.

Wolf, M., De Jong, M., Halldórsson, S. D., Árnason, Ú., & Janke, A. (2022). Genomic impact of whaling in North Atlantic fin whales. Molecular biology and evolution, 39(5), msac094.

Yim, H. S., Cho, Y. S., Guang, X., Kang, S. G., Jeong, J. Y., Cha, S. S., … & Lee, J. H. (2014). Minke whale genome and aquatic adaptation in cetaceans. Nature genetics, 46(1), 88–92.

Yu, J., Nam, B. H., Yoon, J., Kim, E. B., Park, J. Y., Kim, H., & Yoon, S. H. (2017). Tracing the spatio-temporal dynamics of endangered fin whales (*Balaenoptera physalus*) within baleen whale (Mysticeti) lineages: a mitogenomic perspective. Genetica, 145, 603–612.

Zhang, H., Song, L., Wang, X., Cheng, H., Wang, C., Meyer, C. A., … & Li, H. (2021). Fast alignment and preprocessing of chromatin profiles with Chromap. Nature communications, 12(1), 6566.

Zhang, X., Lin, W., Zhou, R., Gui, D., Yu, X., & Wu, Y. (2016). Low major histocompatibility complex class II variation in the endangered Indo-Pacific humpback dolphin (*Sousa chinensis*): Inferences about the role of balancing selection. Journal of Heredity, 107(2), 143–152.

Zhou, C., McCarthy, S. A., & Durbin, R. (2023). YaHS: yet another Hi-C scaffolding tool. Bioinformatics, 39(1), btac808.

