## Supplementary Figures for "One Sea, Different Whales: Genomics Sheds Light on a Small Population of Fin Whales"

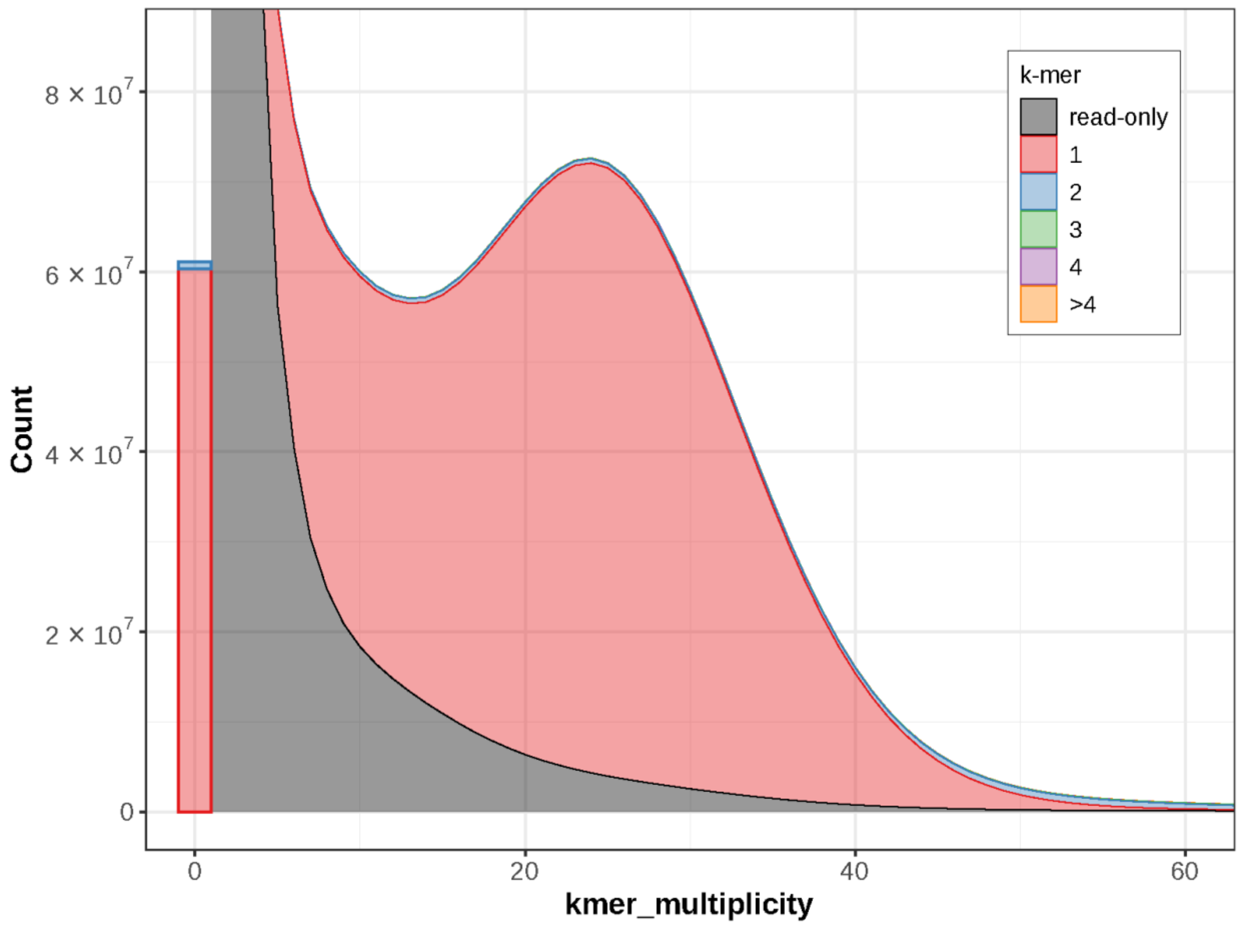


**Figure S1.** Merqury k-mer plots comparing k-mer content of Hi-C raw reads with Bphy_ph2.v2 assembly. The black area of the graphs represents the distribution of k-mers present in the reads but not in the assembly and the red area represents the distribution of k-mers present in the reads and once in the assembly. Other colours show k-mers found multiple times in the genome assembly.

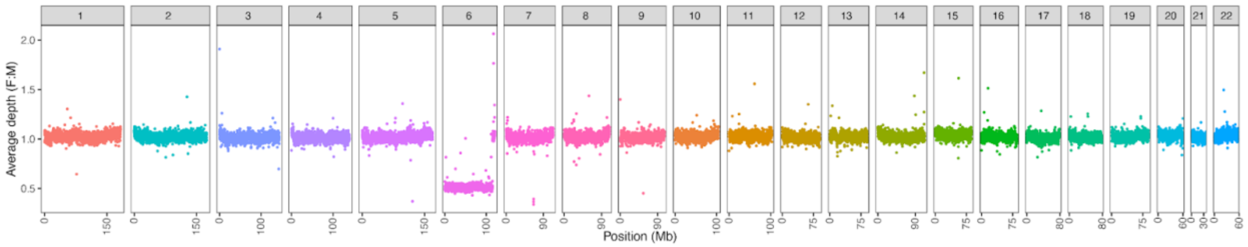


**Figure S2.** Data showing identification of sex chromosomes in *B. physalus* genome assembly, with the sex chromosome displaying 50% lower coverage compared to the autosomes.

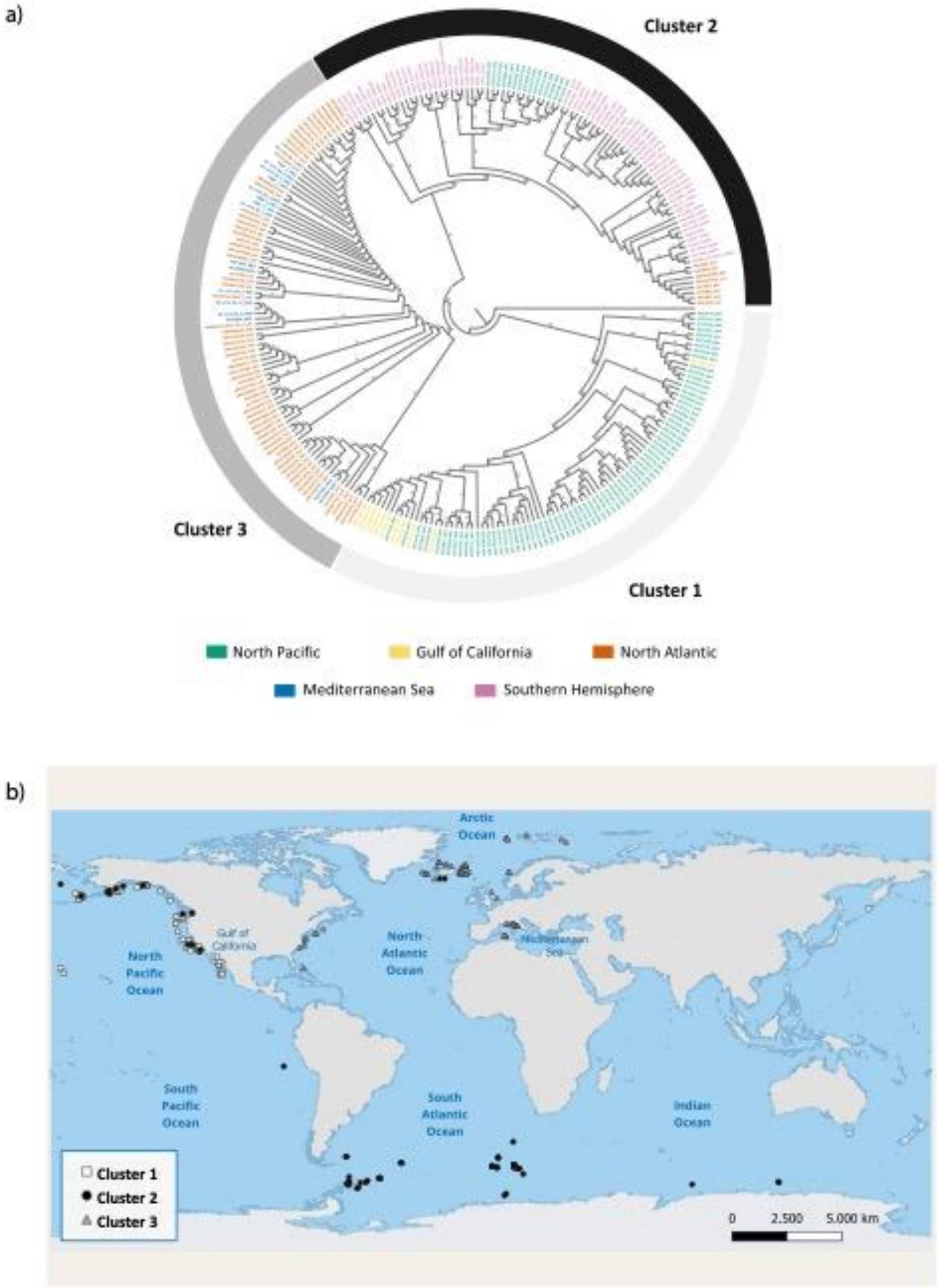


**Figure S3.** Mitochondrial tree. a) Rooted phylogenetic tree based on the maximum likelihood of fin whale mitogenomes used in this study. Colours represent the basins of sample origin: Southern Hemisphere (pink, indicated SHEM), North Pacific (green, indicated NPAC), Gulf of California (yellow, indicated GOC), North Atlantic Ocean (orange, indicated NAT) and Mediterranean Sea (light blue, indicated MED). The numbers indicate bootstrap support values in percentage (0 to 100). The bands outside the tree were used to highlight the presence of three main clusters. The mitogenomic haplotype of *Balaenoptera musculus* (Genbank NC_001601) was used to root the tree. b) Geographic distribution of *B. physalus* samples included in the mitogenomes analysis. Different shapes and colours identify different clusters in the phylogenetic tree. Map produced in QGIS 3.28.8 ([www.qgis.org](http://www.qgis.org))

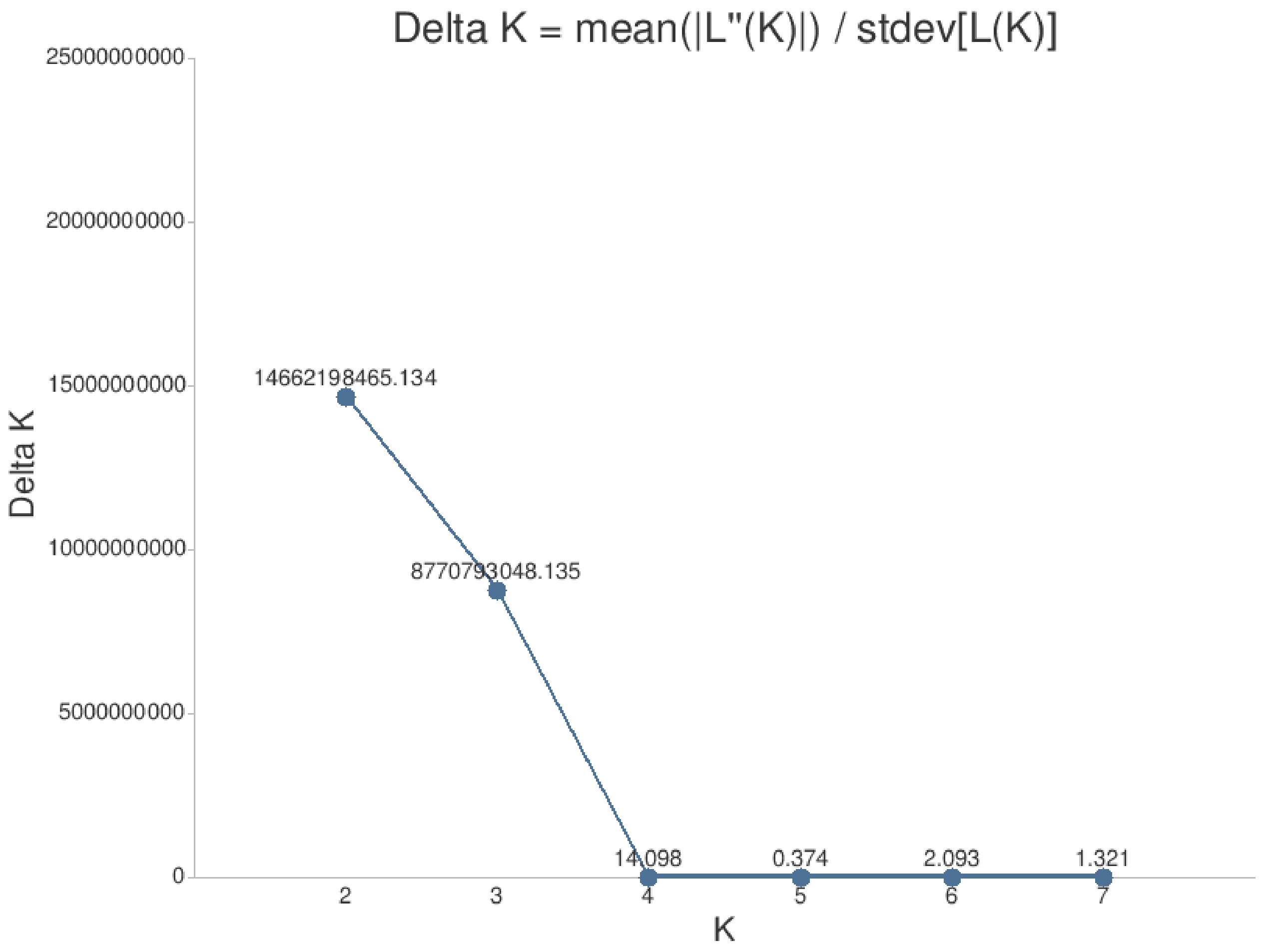


**Figure S4.** Probability of number of clusters (K) using the Delta K method by Evanno et al. (2005).

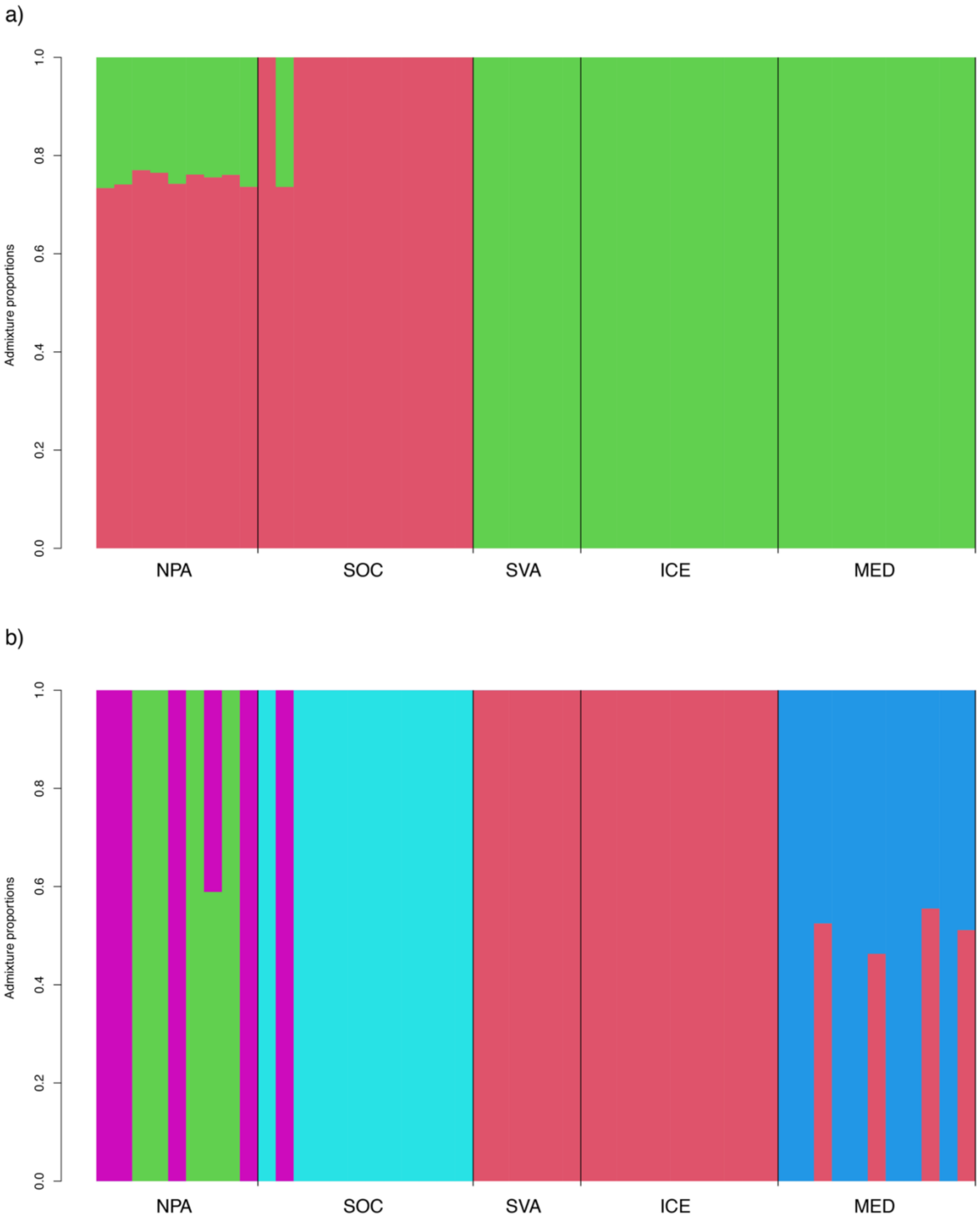


**Figure S5.** Admixture analyses. Each bar in panel d represents an individual, and each colour indicates the proportion of that individual's genome assigned to each of the K clusters. a) K2 and b) K5.

### References

G. Evanno, S. Regnaut, J. Goudet, Detecting the number of clusters of individuals using the software structure: a simulation study, Mol. Ecol. 14 (2005) 2611–2620.
